# Glomerular-Targeted Delivery of Low-Dose Prednisolone Attenuates Established Lupus Nephritis in MRL/lpr Mice

**DOI:** 10.64898/2026.08.01.742194

**Authors:** Kristof Williams, Georgina Agyekum, Akshata Patne, Eleni Markoutsa, David Raj Chellappan, Nathan Hall, Zhi Tian, Nohely Hernandez Soto, Sebastian Cuadrao, Isabella Lozonschi, Liying Fu, Lilliana Haight, Rahul Sharma, Subhra Mohapatra, Lei Wang, Shyam Mohapatra, Ruisheng Liu

**Affiliations:** Department of Molecular Pharmacology and Physiology, Morsani College of Medicine, University of South Florida, Tampa, FL 33602, USA; USF Health Transplant Research Center (UTRC), University of South Florida, Tampa, FL, 33602, USA; Center for Immunity, Inflammation and Regenerative Medicine, Division of Nephrology, University of Virginia School of Medicine, Charlottesville, VA 22903, USA; Department of Molecular Medicine, University of South Florida, Tampa, FL, 33612, USA; Department of Internal Medicine, Institute for Translational Virology and Innovation (ITVI), Morsani College of Medicine, University of South Florida, Tampa, FL 33612, USA; James A Haley VA Hospital, Tampa, FL, 33612, USA; Department of Laboratory Medicine and Pathology, Mayo Clinic Arizona, Scottsdale, AZ 85259, USA

## Abstract

**Background:** Lupus nephritis remains a major cause of chronic kidney disease and kidney failure in systemic lupus erythematosus. Glucocorticoids are central to treatment but are limited by systemic toxicity. We evaluated whether a previously characterized collagen IV α3-targeted liposomal nanoparticle formulation carrying low-dose prednisolone could attenuate established lupus nephritis in MRL/lpr mice.

**Methods:** Female MRL/lpr mice with disease present at treatment initiation and C57BL/6J control mice received saline or collagen IV α3-targeted prednisolone-loaded nanoparticles (Col4-α3-Pred-NPs). Renal outcomes were assessed by longitudinal proteinuria, glomerular filtration rate (GFR), survival, kidney histopathology, renal IgG and C3d deposition, dUTP/TUNEL-associated injury staining, and renal cytokine/chemokine profiling. Body weight, food and water intake, and blood glucose were monitored as measures of general condition and preliminary tolerability.

**Results:** Col4-α3-Pred-NPs improved survival in MRL/lpr mice, reduced cumulative proteinuria burden, and attenuated terminal GFR decline compared with saline-treated MRL/lpr controls. Treatment reduced glomerular and tubulointerstitial injury, lowered composite EGTI histopathology scores, decreased terminal kidney enlargement, reduced glomerular IgG deposition and renal dUTP-positive injury signals, and reduced renal signals for IL-28A/B, IL-7, PD-ECGF, IL-11, CCL6/C10, and IL-15. C3d deposition was not significantly altered. Nanoparticle treatment was not associated with sustained treatment-related increases in blood glucose or body-weight loss during the measured study period.

**Conclusions:** Collagen IV α3-targeted liposomal delivery of low-dose prednisolone attenuated established lupus nephritis in MRL/lpr mice and improved renal structural, functional, inflammatory, and survival outcomes. These findings support further evaluation of glomerulus-targeted nanotherapy as a potential strategy to improve the precision and therapeutic index of glucocorticoid treatment in lupus nephritis.

## Introduction

Systemic lupus erythematosus (SLE) is a chronic systemic autoimmune disease characterized by loss of immunologic self-tolerance and production of autoantibodies [1–5]. These autoantibodies form circulating immune complexes that deposit in multiple tissues, triggering complement activation, inflammatory-cell recruitment, and progressive organ damage [6–10]. SLE is clinically heterogeneous and disproportionately affects women, with a female-to-male ratio of approximately 6:1 [2–4]. Lupus nephritis (LN), one of the most severe organ manifestations of SLE, occurs in approximately 40-60% of patients and remains a major contributor to chronic kidney disease and progression to end-stage kidney disease (ESKD) [6–8,11,12].

The pathogenesis of LN is driven by immune-complex-mediated injury within the renal microvasculature [6–10]. Deposition of immune complexes in glomerular capillaries, the glomerular basement membrane (GBM), and the mesangium initiates complement activation, endothelial dysfunction, and inflammatory-leukocyte recruitment [6–10,13,14]. These processes promote progressive glomerular damage, tubulointerstitial injury, fibrosis, and impaired renal function [6–9]. Despite advances in immunomodulatory therapy, renal flares remain common and sustained remission is difficult to achieve, underscoring the need for more effective and targeted approaches [11,12,15–20].

Current LN treatment relies heavily on systemic immunosuppression [11,12,15,16,20]. Glucocorticoids such as prednisone or prednisolone remain central to initial therapy because of their potent anti-inflammatory and immunosuppressive effects [11,12,15,16,25]. However, their lack of tissue specificity results in off-target effects that compromise long-term treatment sustainability [25–28]. Prolonged systemic exposure is associated with osteoporosis, avascular necrosis, metabolic dysregulation, hepatic steatosis, electrolyte imbalance, and cumulative organ damage [25–28]. These adverse effects constrain dosing strategies and contribute substantially to long-term morbidity in patients with SLE [25–28].

An unmet need in LN management is the development of therapies that retain anti-inflammatory activity while limiting systemic exposure [11,12,15,16,25–28]. Nanotechnology-based drug-delivery systems offer a potential route to organ-directed treatment [29,30]. Liposomes are widely used as nanoparticle (NP) carriers because they can encapsulate therapeutic agents, but conventional formulations generally lack tissue specificity [29,30]. Targeted NP platforms have therefore been engineered to enhance renal or glomerular accumulation through particle size, surface charge, and ligand conjugation [29,30].

One such strategy exploits the structural and biological properties of collagen IV within the GBM. The α3 chain of collagen IV (Col4-α3) is a key GBM component that is accessible to circulating molecules through the fenestrated glomerular endothelium [31–35]. Thus, Col4-α3 is a biologically rational target for site-directed delivery [29–35]. Building on this rationale, we previously developed a glomerulus-targeted liposomal NP system by conjugating a Col4-α3-binding peptide to prednisolone-loaded liposomes [35]. The formulation exhibited high-affinity binding, sustained drug release for 24–48 hours, approximately twofold greater kidney fluorescence than non-targeted NPs, and detectable uptake in more than 80% of glomeruli [35]. The platform also produced therapeutic benefit when administered during an earlier disease window at a dose described as approximately 20-fold lower than doses used in the cited preclinical literature [22–24, 35]. However, clinical treatment begins after disease onset; therefore, the translational relevance of this approach depends on efficacy after renal disease has developed. In the present study, we evaluated Col4-α3-targeted prednisolone-loaded nanoparticles (Col4-α3-Pred-NPs) in MRL/lpr mice treated after lupus-associated renal disease had begun. We hypothesized that glomerulus-directed delivery would attenuate renal inflammation, reduce immune-complex-associated injury, and preserve kidney function. We also assessed general condition and selected metabolic and body-composition measures as preliminary indicators of tolerability; the study was not designed to establish reduced systemic exposure relative to free prednisolone.

## METHODS

### Synthesis and Characterization of Liposomal Nanoparticles

Col4-α3-Pred-NPs were prepared as previously described [35]. Briefly, prednisolone was encapsulated within PEGylated liposomes, and a collagen IV α3-binding peptide (KLWVLPKGGGC) was conjugated to maleimide-functionalized groups on the liposomal surface through thiol–maleimide chemistry. The resulting liposomes were approximately 100 nm in diameter, exhibited a peptide-conjugation efficiency of approximately 75%, and contained an estimated 1,000 targeting peptides per liposome. The formulation used in the present study was prepared according to the previously reported procedure without modification.

### Animals and Ethical Approval

All animal experiments were conducted in accordance with the National Institutes of Health Guide for the Care and Use of Laboratory Animals and were approved by the Institutional Animal Care and Use Committee at the University of South Florida (IACUC protocol no. 11754). Mice were obtained from The Jackson Laboratory and acclimated for 3 weeks before experimental procedures. Animals were housed under specific pathogen-free conditions at a controlled temperature of 22 ± 2°C, with controlled humidity, five mice per cage, a 12-hour light/dark cycle, and ad libitum access to standard chow and water. Cages were randomly assigned to experimental groups.

Animals were monitored regularly for health status, body weight, and clinical signs of disease progression. Food and water intake were recorded per cage. Humane endpoints were predefined and enforced throughout the study.

### Animal Models Study Design

MRL/MpJ-Faslpr/J mice were used to evaluate therapeutic efficacy, and C57BL/6J (B6) mice were included as non-lupus controls for preliminary tolerability assessment [36–41]. Treatment began at 12 weeks of age. Four groups were studied: B6 saline, B6 Col4-α3-Pred-NP, MRL/lpr saline, and MRL/lpr Col4-α3-Pred-NP. Col4-α3-Pred-NP was administered twice weekly by retro-orbital sinus injection at the reported prednisolone-equivalent dose of 0.34 mg/kg/day. Retro-orbital administration was selected to provide consistent systemic delivery. Only female mice were included because LN develops earlier and more consistently in female than in male MRL/lpr mice [36–41]. Renal function and injury were assessed as described below.

### Proteinuria Assessment

Urine samples were collected weekly by spot collection at approximately 9:00 AM from non-fasted mice, with approximately 200 µL collected per sample. Samples were centrifuged at 400 × g for 5 minutes and stored at −80°C until analysis. Urinary protein concentration was quantified using the Pierce™ BCA Protein Assay Kit (Thermo Fisher Scientific). Urinary creatinine was measured by high-performance liquid chromatography at the O’Brien Center Biomedical Resource Core, University of Alabama at Birmingham. Urinary protein-to-creatinine ratios were calculated to assess renal injury and disease progression.

### Blood Glucose Assessment

Blood glucose was measured longitudinally from plasma samples collected at baseline and subsequent study time points. After blood collection, plasma was separated and used for glucose measurement with a handheld glucose meter and compatible test strips. Glucose concentrations were recorded in mg/dL.

### Glomerular Filtration Rate (GFR) Measurement

Glomerular filtration rate (GFR) was assessed biweekly by transdermal FITC-sinistrin clearance. Mice received FITC-sinistrin prepared at 28 mg/mL by retro-orbital injection at 7 mg/100 g body weight, and clearance was recorded in conscious mice using the MediBeacon Preclinical MX Transdermal GFR Monitor. Data was analyzed using MediBeacon Studio software (version 2.3).

### Survival Analysis

Animal survival was monitored throughout the study period. Deaths attributable to experimental handling or procedural complications were censored and excluded from survival event analysis. Survival curves were generated using the Kaplan–Meier method, and statistical significance was assessed using the log-rank (Mantel-Cox) test. All survival analyses were performed using GraphPad Prism (version 10, GraphPad Software, San Diego, CA, USA).

### Tissue Collection and Processing

At study termination or upon reaching predefined humane endpoints, animals were anesthetized with 2% isoflurane and perfused with physiological saline to remove circulating blood. The kidneys, spleen, liver, heart, brain, and lymph nodes were harvested, weighed, and processed for downstream analyses. Kidneys designated for histology were fixed in 4% paraformaldehyde (PFA) prior to tissue processing.

### Histology and Pathological Scoring

Tissue samples were fixed in 4% PFA for 24 hours, then immersed in 70% ethanol for an additional 24 hours. Paraffin embedding, sectioning, and periodic acid–Schiff (PAS) staining was performed at the H. Lee Moffitt Cancer Center & Research Institute. Hematoxylin and Eosin (H&E) staining was performed on 4µm-thick tissue sections using a standard H&E protocol. Whole tissue section images were acquired at magnifications ranging from 4× to 40× using an Olympus VS120 slide scanning system (Olympus, Tokyo, Japan), at the Lisa Muma Weitz Imaging Core Facility, University of South Florida (Tampa, FL). Renal injury was assessed using an adapted, semiquantitative EGTI scoring framework, which evaluates glomerular, tubulointerstitial, endothelial, and tubular pathology [32]. Scoring was performed in a blinded manner by a board-certified pathologist, and 75 sections from each animal were evaluated. Composite injury scores were calculated by averaging scores across all analyzed images and tissue sections from each animal.

### Immunofluorescence Staining

Paraffin-embedded tissue sections were deparaffinized, permeabilized, and blocked prior to immunofluorescence staining. Endogenous IgG deposition was detected using Alexa Fluor 488–conjugated donkey anti-mouse IgG (Abcam, Cambridge, UK; 1:1000). Complement component C3d was detected using a goat anti-C3d antibody (R&D Systems, Minneapolis, USA; 10 µg/mL), followed by incubation Alexa Fluor 594 – conjugated donkey anti-goat secondary (Abcam, USA; 1:800). Nuclei were counterstained with DAPI mounting media. All sections were imaged using identical acquisition settings to allow direct comparison of fluorescence intensity and distribution among experimental groups.

### TUNEL/dUTP-Associated Injury Staining

DNA fragmentation was assessed in paraffin-embedded kidney sections using the Click-iT™ Plus TUNEL Assay for In Situ Apoptosis Detection with Alexa Fluor™ 594 dye (Invitrogen, Thermo Fisher Scientific; C10618) according to the manufacturer’s instructions. Nuclei were counterstained with DAPI, and sections were imaged using identical acquisition settings across experimental groups.

### Cytokine and Chemokine Profiling

Inflammatory mediators were assessed using membrane-based cytokine arrays. Forty analytes were assessed in B6 samples and 111 analytes in MRL/lpr samples (Proteome Profiler Mouse Cytokine Array Kits ARY006 and ARY028; R&D Systems, Minneapolis, MN, USA). Kidney homogenates were processed according to the manufacturer’s protocol. Chemiluminescent signals were acquired using a Bio-Rad ChemiDoc system and quantified by densitometry. Duplicate spot intensities were summarized and normalized to the internal positive-reference spots on each membrane before group comparison. All array analytes were included in the initial analysis.

### DEXA and Body-Composition Analysis

In a separate pilot cohort presented in Supplementary Figure 2, bone mineral content, bone mineral density, and body fat percentage were assessed longitudinally in MRL/lpr mice at 6, 13, and 15 weeks of age using an Insight Vet DXA system. The 6-week measurement was obtained before treatment initiation and served as the baseline. Mice subsequently received blank nanoparticles, prednisolone-loaded Col4-α3–targeted nanoparticles, or free prednisolone twice weekly at the reported average daily-equivalent dose of 0.34 mg/kg/day through 15 weeks of age. Mice were anesthetized with 2% isoflurane, positioned prone with the limbs extended, and scanned according to the manufacturer’s instructions. Bone mineral content, bone mineral density, and body fat percentage were recorded. These exploratory measurements were conducted in the pilot cohort and were not part of the primary efficacy study.

### Statistical Analysis

All statistical analyses were performed using GraphPad Prism version 10 (GraphPad Software, San Diego, CA, USA). Data are presented as mean ± standard error of the mean (SEM). Normality was assessed before selecting parametric or nonparametric statistical tests. Longitudinal data were analyzed using repeated-measures analysis of variance (ANOVA) where appropriate. Two-group comparisons were performed using unpaired Student’s t-tests or the corresponding non-parametric tests when normality assumptions of normality were not met. Survival outcomes were analyzed using the Kaplan-Meier method and compared using the log-rank (Mantel-Cox) test. A two-sided p-value < 0.05 was considered statistically significant.

## RESULTS

### Col4-α3–targeted prednisolone nanoparticles improve survival without sustained hyperglycemia or body-weight loss

Kaplan–Meier analysis showed a significant difference in survival distributions across the four experimental groups (log-rank p = 0.0013; Figure 1A). Median survival was reached only in saline-treated LPR mice, whereas 80% of NP-treated LPR mice survived to the study endpoint (Table 1). B6 mice maintained high survival irrespective of treatment (Figure 1A).

**Figure 1.**
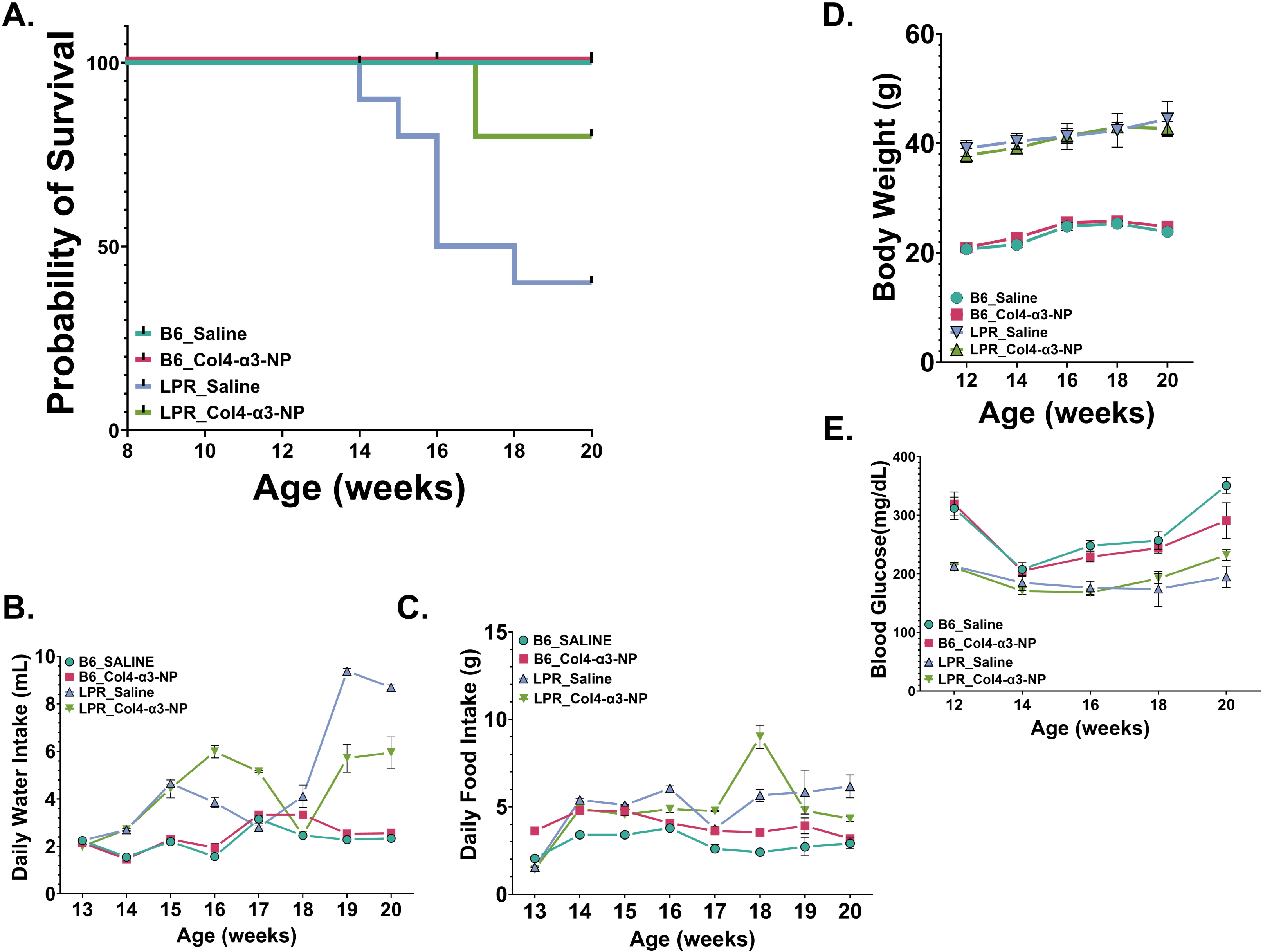
Survival and general condition of B6 and MRL/lpr mice. **A.** Kaplan–Meier survival curves for C57BL/6J (B6) and MRL/lpr mice treated with saline or collagen IV α3–targeted prednisolone-loaded nanoparticles (Col4-α3-Pred-NPs). Tick marks indicate censored observations. Survival distributions were compared using log-rank (Mantel–Cox) test (χ² = 15.74, df = 3, p = 0.0013). **B.** Longitudinal water intake in B6 and MRL/lpr mice. Treatment-associated differences were observed in both strains, with diverge between MRL/lpr treatment groups at later time points. **C.** Longitudinal food intake in B6 and MRL/lpr mice. B6 mice treated with Col4-α3-Pred-NPs exhibited higher food intake than salin treated B6 controls, whereas MRL/lpr mice showed no significant overall treatment effect. **D.** Longitudinal body weight in B6 and MRL/lpr mice. Body weight changed significantly over time for all groups, but there was no significant treatment effect or time × treatment interaction. **E.** Longitudinal glucose measurements in non-fasted B6 and MRL/lpr mice. Glucose varied over time within each strain, but no significant treatment effect was detected. Data are presented as mean ± SEM. Longitudinal outcomes were analyzed using the statistical models specified in the Methods *The number of animals contributing data at each longitudinal time point varied because of mortality, censoring, and sam availability; animal-level disposition and availability by study week are provided in Supplementary Table S1.

**Table 1:** Survival and censoring summary for B6 and MRL/lpr mice treated with saline or Col4-α3-Pred-NPs.

| Group | n | Death Events | Early Censored | Reached Week 20 |
| --- | --- | --- | --- | --- |
| B6_Saline | 10 | 0 | 3 | 7 |
| B6_Col4-α3-NP | 10 | 0 | 1 | 9 |
| LPR_Saline | 10 | 6 | 0 | 4 |
| LPR_Col4-α3-NP | 10 | 2 | 0 | 8 |

**Note:** B6, C57BL/6J; MRL/lpr, MRL/MpJ-Fas<sup>lpr</sup>/J lupus-prone mice; Col4-α3-NP, collagen IV α3-targeted prednisolone-loaded NPs. Death events include animals that died during the study period and were included as survival events. Early-censored animals were excluded from event analysis because death or removal was attributed to procedural or non-disease-related causes. Survival was analyzed using Kaplan–Meier curves and log-rank Mantel–Cox test. $n = 10$ mice/group at enrollment.

Food and water intake, body weight, and blood glucose were monitored longitudinally as measures of general condition. In B6 mice, NP-treated animals showed higher water intake than saline-treated controls, with significant effects of time, treatment, and time × treatment interaction (ANOVA, p < 0.0002; Figure 1B). In lupus-prone MRL-LPR mice, water intake increased with age and diverged between treatment groups at later time points, with NP-treated mice maintaining higher intake than saline-treated controls (ANOVA, p < 0.0004; Figure 1B)

Food intake differed between B6 treatment groups, with NP-treated mice maintaining higher intake than saline-treated controls (ANOVA, p < 0.009; Figure 1C). Food intake varied over time but showed no consistent treatment-associated separation between saline- and NP-treated LPR mice (ANOVA, p = 0.4; Figure 1C). Body weight increased progressively from 12 to 20 weeks of age in LPR and B6 mice, with substantial overlap between treatment groups at all time points (Figure 1D). ANOVA demonstrated a significant effect of time (p=0.0174), but no main effect of treatment or time × treatment interaction (p > 0.5), indicating that NP-delivered prednisolone did not significantly alter body-weight trajectories.

Blood glucose was measured longitudinally to assess whether NP-delivered prednisolone produced systemic glucocorticoid-associated metabolic effects. (Figure 1E) Values were significantly higher in B6 mice when compared to LPR mice over time (ANOVA, p < 0.0034). When comparing B6 groups, glucose levels changed significantly over time (p < 0.0001), but there was no significant effect of treatment (p > 0.1) Similarly, between LPR groups, glucose levels varied over time (p = 0.0002), but NP treatment did not significantly alter glucose compared with saline-treated controls (p > 0.2)

### Glomerular targeted prednisolone therapy reduces proteinuria burden and attenuate terminal GFR decline

Renal function was assessed longitudinally using urinary protein excretion and glomerular filtration rate (GFR). Saline-treated MRL/lpr mice exhibited a greater proteinuria burden over the study period, with proteinuria increasing through the later stages of disease, whereas Col4-α3-Pred-NP–treated mice showed lower and more stable proteinuria values (p < 0.05; Figure 2A). Summary analyses further demonstrated a lower overall proteinuria burden in treated mice, including reduced area under the curve (AUC), baseline-normalized AUC, and peak proteinuria (p < 0.02; Figure 2B).

**Figure 2.**
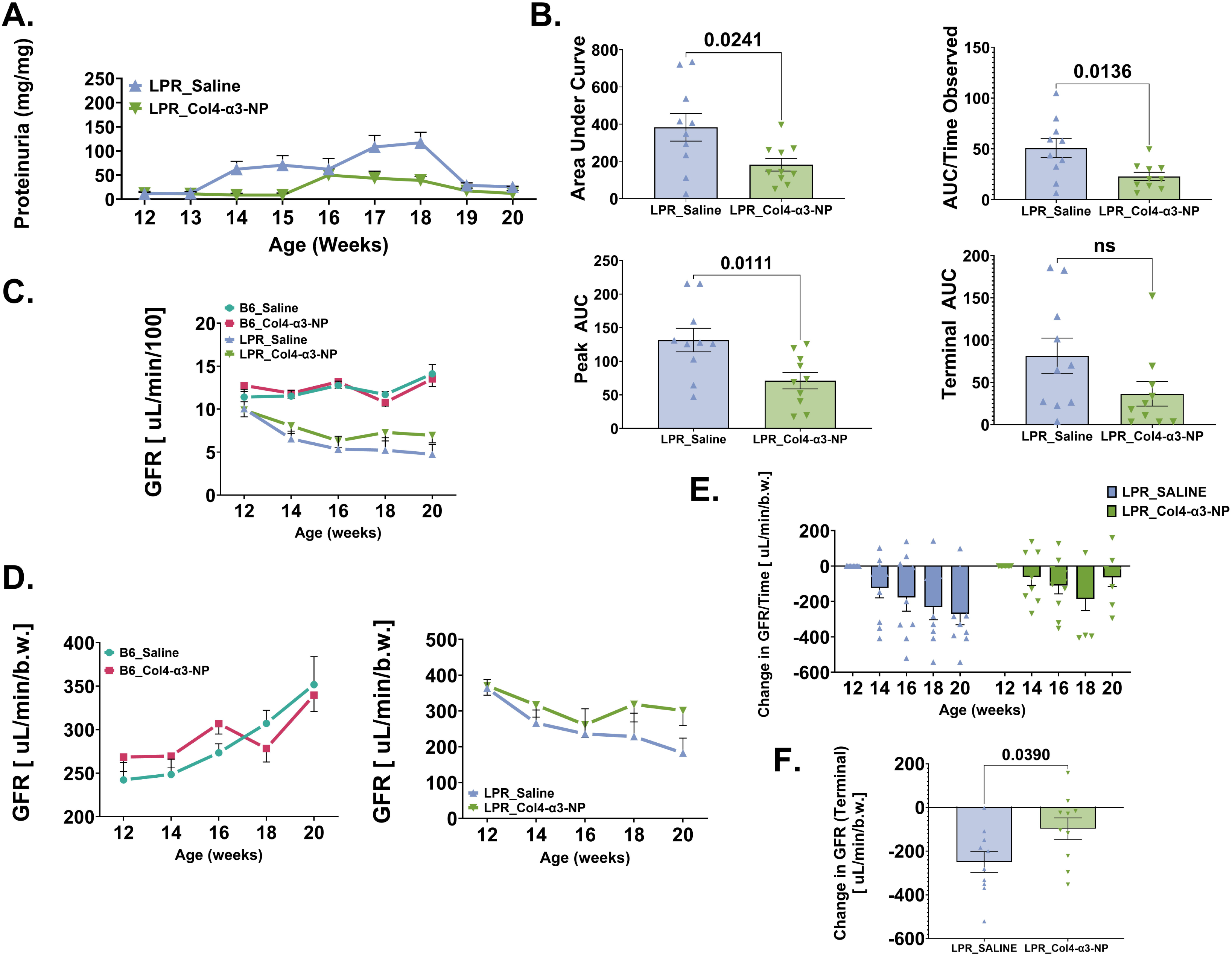
Col4-α3-Pred-NPs reduce proteinuria burden and attenuate terminal GFR decline in MRL/lpr mice. A. Longitudinal urinary protein-to-creatinine ratios in MRL/lpr mice treated with saline or collagen IV α3–targeted prednisolone-loaded nanoparticles (Col4-α3-Pred-NPs). **B.** Proteinuria summary measures, including area under the curve (AUC), baseline-normalized AUC, peak proteinuria, and terminal proteinuria. **C.** Absolute longitudinal glomerular filtration rate (GFR) in saline- and Col4-α3-Pred-NP–treated B6 and MRL/lpr mice. **D.** Body-weight-normalized longitudinal GFR displayed separately for B6 mice (left) and MRL/lpr mice (right). **E.** Change in GFR from baseline in saline- and Col4-α3-Pred-NP–treated MRL/lpr mice. **F.** Terminal change in GFR from baseline in MRL/lpr mice. Data are presented as mean ± SEM, with individual animals shown where applicable. Proteinuria summary measures and terminal GFR change were analyzed using unpaired two-tailed t tests. Body-weight-normalized longitudinal GFR was analyzed using mixed-effects models fitted by restricted maximum likelihood with Geisser–Greenhouse correction. Change-from-baseline GFR was analyzed using two-way ANOVA. n = 10 mice/group at enrollment. *The number of animals contributing data at each longitudinal time point varied because of mortality, censoring, and sample availability; animal-level disposition and availability by study week are provided in Supplementary Table S1.

Terminal proteinuria was also lower in treated mice but did not reach statistical significance (p = 0.09). Together, these findings indicate that Col4-α3-Pred-NP treatment attenuated the cumulative proteinuria burden in lupus-prone mice.

GFR was evaluated longitudinally using both absolute and body-weight-normalized measurements. Absolute GFR values for all four experimental groups are shown in Figure 2C. Body-weight-normalized GFR values are shown separately for B6 and MRL/lpr mice in Figure 2D and were analyzed using within-strain mixed-effects models. In B6 mice, normalized GFR changed significantly over time (p = 0.0013), but there was no significant treatment effect (p = 0.5610) or time × treatment interaction (p = 0.3390), indicating no detectable treatment-associated alteration in renal filtration. In MRL/lpr mice, normalized GFR also changed significantly over time (p = 0.0402). The treatment effect approached but did not reach statistical significance (p = 0.0599), and the time × treatment interaction was not significant (p = 0.5826). Although NP-treated MRL/lpr mice exhibited numerically higher normalized GFR values at later time points, the overall longitudinal trajectories were not significantly different between groups.

Change-from-baseline GFR was further evaluated in MRL/lpr mice. Two-way ANOVA demonstrated significant effects of time (p = 0.0019) and treatment (p = 0.0252), but no significant time × treatment interaction (p = 0.3842; Figure 2E). Across the evaluated time points, Col4-α3-Pred-NP–treated mice exhibited a smaller mean reduction in GFR from baseline than saline-treated controls. The terminal decline in GFR from baseline was also significantly smaller in NP-treated mice than in saline-treated MRL/lpr controls (p = 0.039; Figure 2F). These secondary findings support attenuation of GFR decline, although the nonsignificant interaction indicates that the shapes of the longitudinal trajectories were not statistically different.

### Glomerular targeted prednisolone therapy ameliorates renal structural injury and pathological severity in lupus nephritis

Renal histopathology was evaluated to determine whether functional improvements were associated with structural protection. Representative H&E-stained kidney sections demonstrated preserved renal architecture in B6 mice regardless of treatment (Figure 3A). In contrast, saline-treated LPR mice exhibited pronounced pathological features, including interstitial inflammatory infiltrates, acute tubular injury with loss of brush borders, and intratubular casts. NP-treated MRL/lpr mice showed partial attenuation of these pathological features compared with untreated controls. Periodic acid–Schiff (PAS) staining further revealed intact glomerular architecture in kidney sections from B6 mice, characterized by delicate glomerular basement membranes, minimal mesangial expansion, and preserved capillary loops (Figure 3B). PAS-stained renal sections from Saline-treated MRL/lpr mice displayed marked glomerular abnormalities, including basement membrane thickening and reduplication, mesangial proliferation, and glomerulosclerosis. These structural abnormalities were reduced in nanoparticle-treated MRL/lpr mice, indicating amelioration of glomerular injury.

**Figure 3.**
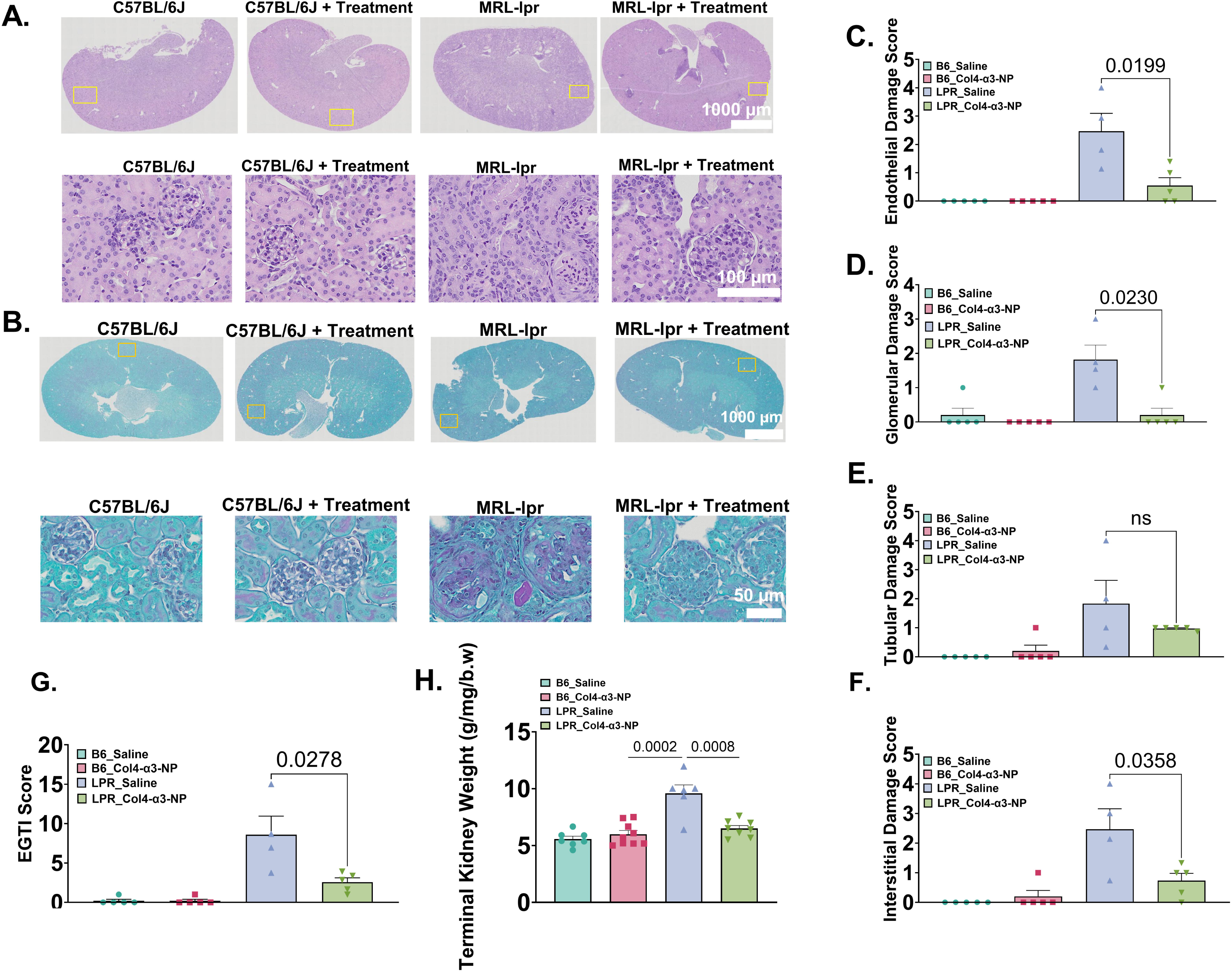
Histopathologic assessment of renal injury in B6 and MRL/lpr mice. **A–B**. Representative kidney sections stained with hematoxylin and eosin (H&E) and periodic acid–Schiff (PAS). B6 kidneys show preserved renal architecture, whereas saline-treated MRL/lpr kidneys exhibited prominent glomerular and tubulointerstitial injury. Renal lesions were attenuated in Col4-α3-Pred-NP–treated MRL/lpr mice. **C–G**. Semiquantitative endothelial, glomerular, tubular, and interstitial injury (EGTI) component scores and composite EGTI score Minimal injury was observed in B6 groups, whereas Col4-α3-Pred-NP treatment reduced several measures of renal injury in MRL/ mice. **H**. Terminal kidney weight in B6 and MRL/lpr mice treated with saline or Col4-α3-Pred-NPs. Data are presented as mean ± SEM individual animals shown. Exact p values are displayed in the figure; p < 0.05, *p < 0.01, **p < 0.001; ns, not significant. Scale bars are shown in the images.

Quantitative assessment using the EGTI scoring system demonstrated minimal pathology across all compartments in B6 groups (ANOVA, p > 0.05; Figures 3C-G). In MRL/lpr mice, NP-delivered prednisolone treatment significantly reduced glomerular and tubulointerstitial injury scores and lowered the composite EGTI score, with a trend toward reduced tubular injury. (ANOVA, p < 0.003; Figures 3C-G) Terminal kidney weight, normalized to body weight, was significantly increased in saline-treated MRL/lpr mice compared with B6 controls, consistent with renal inflammation and hypertrophy (Unpaired t-test, p = 0.0002; Figure 3H). Collectively, these findings demonstrate that Col4-α3-targeted NP delivery of prednisolone mitigates both glomerular and tubulointerstitial injury and pathology in lupus-prone mice.

### Glomerular targeted prednisolone therapy reduces renal injury signals and IgG deposition but not C3d deposition

Immunofluorescence analysis demonstrated substantial glomerular IgG-associated fluorescence in saline-treated LPR mice and visibly lower glomerular IgG signal in Col4-α3-Pred-NP–treated mice. Quantitative analysis confirmed significantly lower relative IgG fluorescence intensity in the treated group (unpaired two-tailed t test, p < 0.01; n = 5; Figure 4A, C). This finding is consistent with reduced glomerular IgG accumulation but does not establish reduced circulating autoantibody production.

**Figure 4.**
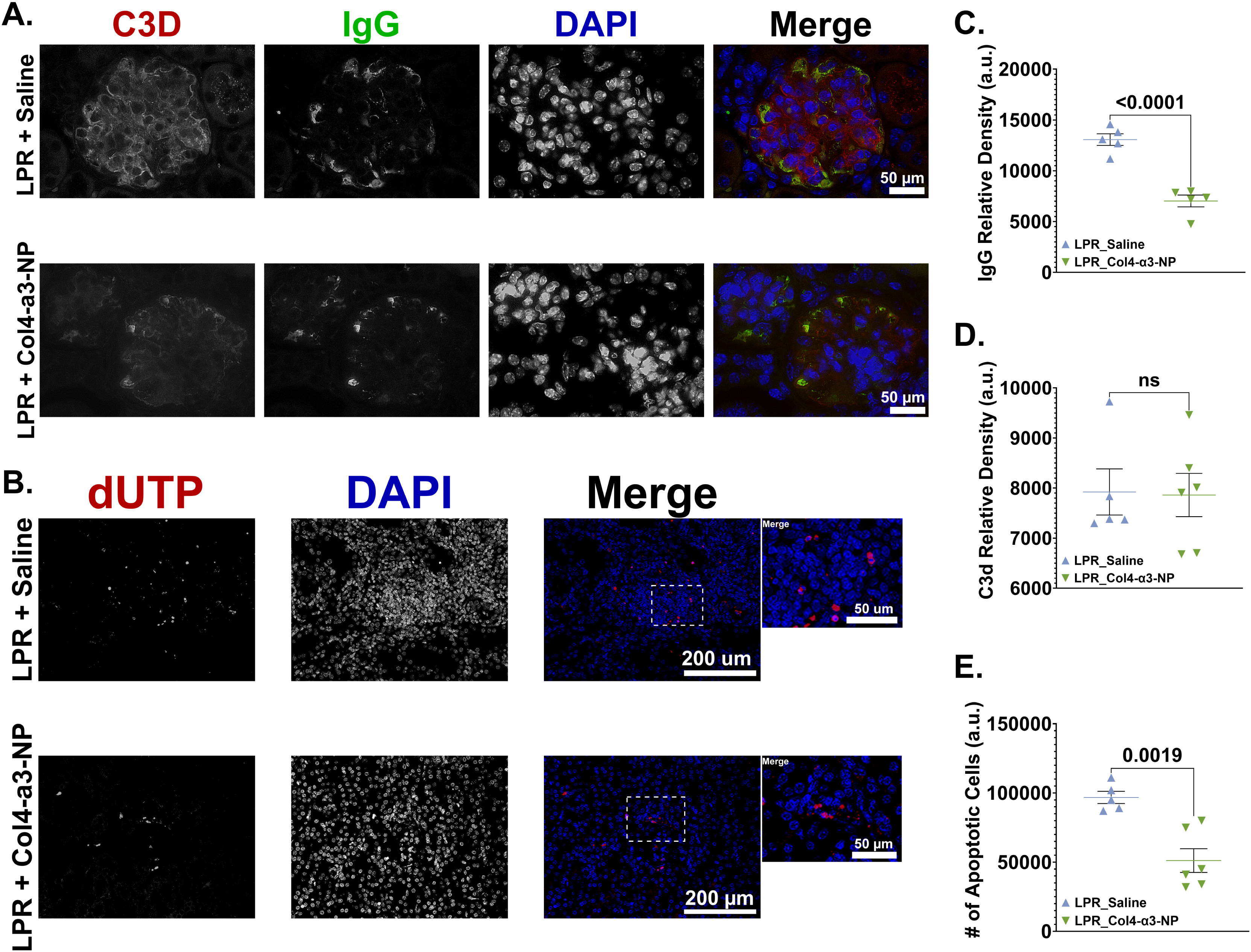
Col4-α3-Pred-NP treatment reduces glomerular IgG fluorescence and DNA-fragmentation–associated renal injur in MRL/lpr mice. **A.** Representative immunofluorescence images of renal IgG-associated fluorescence (green) with DAPI nuclear counterstaining (blue) in saline- and Col4-α3-Pred-NP–treated MRL/lpr mice. **B.** Representative renal sections stained for TUNEL/dUTP-positive DNA-fragmentation signal (red) and DAPI (blue). Insets show higher-magnification views of selected regions. **C.** Quantification of relative glomerular IgG fluorescence intensity. **D.** Quantification of relative glomerular C3d fluorescence intensity. **E.** Quantification of TUNEL/dUTP-positive fluorescence signal. Data are presented as mean ± SEM with individual animals shown. Exact p values are displayed in the figure, p < 0.05; ns, not significant.

Glomerular C3d fluorescence did not differ between saline- and NP-treated MRL/lpr mice (p = 0.9251; Figure 4D), indicating that the reduction in IgG-associated fluorescence was not accompanied by a detectable change in terminal C3d deposition. TUNEL/dUTP staining was used to assess DNA-fragmentation–associated tissue injury. Saline-treated LPR kidneys exhibited widespread TUNEL/dUTP-positive staining, whereas treated kidneys showed lower staining intensity and fewer positive foci (p = 0.0019; Figure 4B, E). Together, these findings indicate that Col4-α3-Pred-NP treatment was associated with reduced glomerular IgG fluorescence and lower DNA-fragmentation–associated renal injury signals, without a detectable change in C3d deposition.

### Glomerular targeted prednisolone therapy suppresses renal inflammatory mediator expression

Renal cytokine and chemokine signals were profiled in LPR kidney homogenates using a membrane-based array. A heatmap of the most treatment-responsive mediators showed lower signals in Col4-α3-Pred-NP-treated samples than in saline-treated samples (Figure 5A). Because the heatmap was generated from treatment-responsive analytes, the apparent group separation should be interpreted descriptively rather than as independent validation.

**Figure 5.**
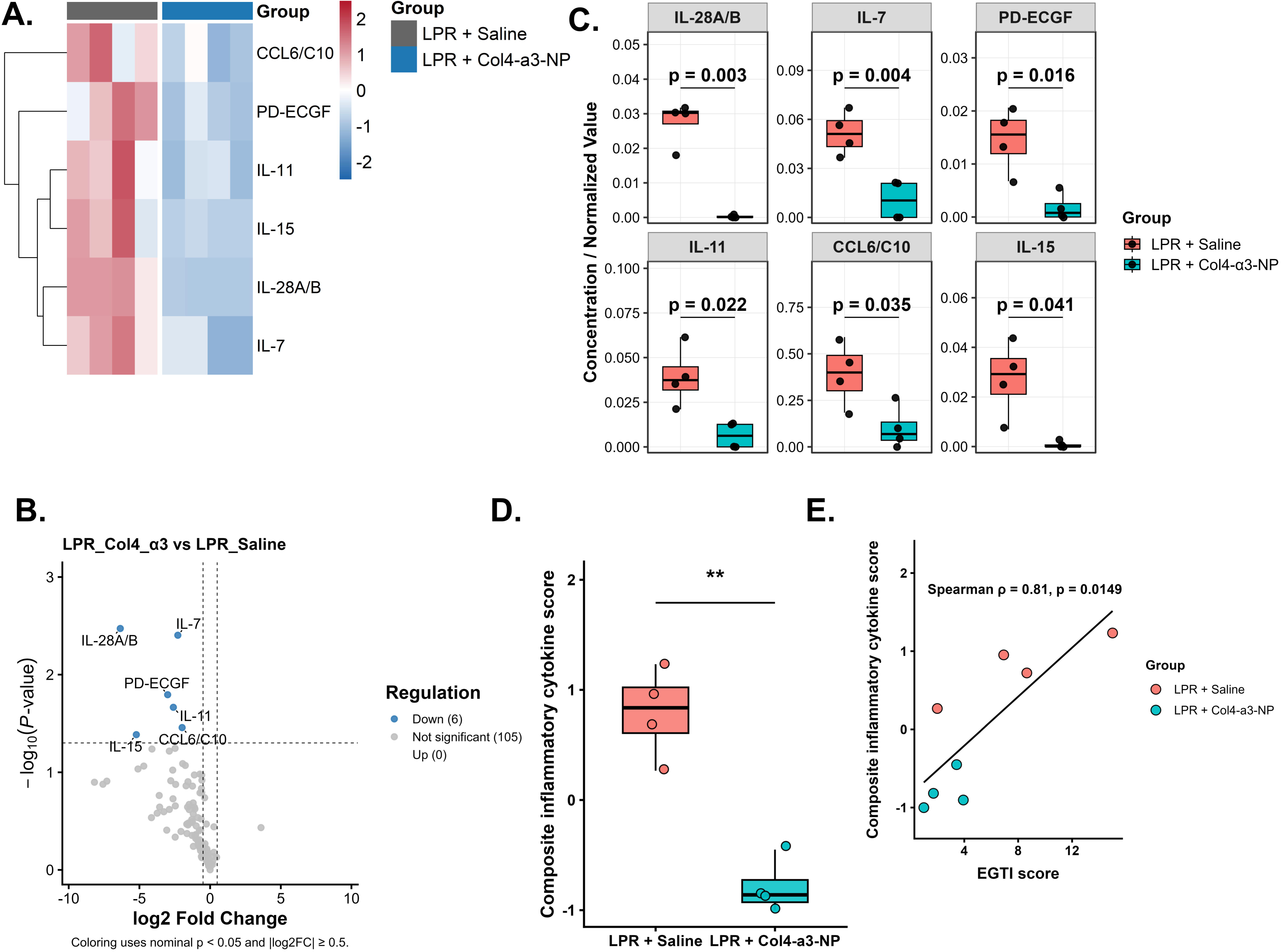
Col4-α3-Pred-NP treatment is associated with reduced renal inflammatory mediator levels in MRL/lpr mice. **A.** Heatmap of renal cytokines and chemokines meeting the prespecified nominal significance and fold-change thresholds in Col4-α3-Pred-NP–treated and saline-treated MRL/lpr mice. **B.** Relative densitometric measurements of IL-28A/B, IL-7, PD-ECGF, IL-11, CCL6/C10, and IL-15. **C.** Volcano plot of renal cytokine-array measurements comparing treatment groups. Six mediators met the nominal thresholds of *p* < 0.05 and absolute log₂ fold change ≥ 0.5; no mediators meeting these criteria were increased in the treated group. **D.** Composite inflammatory mediator score calculated from the six mediators identified in the array analysis. **E.** Spearman correlation between the composite inflammatory mediator score and the composite EGTI histological injury score. Data are shown as individual biological replicates with group summaries. Individual mediators were compared using multiple unpaired *t* tests, the composite score was compared using an unpaired two-tailed *t* test, and its association with EGTI score was assessed using Spearman correlation. Nominal *p* values are shown; *p* < 0.05, \**p* < 0.01. *n* = 4 mice/group.

Using nominal p < 0.05 and |log2 fold change| ≥ 0.5, six mediators showed lower signals in treated kidneys, and no analytes met the corresponding threshold for increased expression (Figure 5B,C). The six mediators were IL-28A/B, IL-7, PD-ECGF, IL-11, CCL6/C10, and IL-15. Individual densitometric comparisons and the six-marker composite score were lower in treated mice (Figure 5C,D). The composite score correlated positively with EGTI score (Spearman ρ = 0.81, p = 0.0149; Figure 5E).

These exploratory findings are consistent with reduced renal inflammatory-mediator abundance but require false-discovery-rate correction and independent validation before individual cytokines are interpreted as confirmed treatment targets.

## Discussion

In the present study, Col4-α3-targeted nanoparticle-mediated prednisolone delivery significantly attenuated established LN in MRL/lpr mice. Treatment improved survival, reduced proteinuria burden, attenuated terminal GFR decline, decreased renal histopathological injury and glomerular immune-complex deposition, and suppressed renal inflammatory mediator expression without overt treatment-related adverse effects. Collectively, these findings support glomerulus-targeted nanotherapy as a strategy with the potential to improve the therapeutic index of glucocorticoids in LN [11,12,15,16,25–30].

A key strength of this study is the treatment of established LN rather than prophylactic intervention, as previously reported [35]. MRL/lpr mice were treated during active immune-complex deposition and progressive renal dysfunction, more closely reflecting the stage at which patients typically begin therapy [36–41]. The attenuation of disease progression under these conditions suggests that Col4-α3-targeted nanoparticle therapy remains effective after pathogenic renal inflammation has been initiated. The collagen IV α3-binding peptide also provides a biologically rational targeting strategy because its GBM target is accessible to circulating NPs through the fenestrated glomerular endothelium [31–34]. By leveraging this accessibility, the platform was designed to increase renal prednisolone accumulation while limiting systemic exposure.

Among the functional outcomes, the most robust treatment effect was the substantial reduction in proteinuria in NP-treated lupus-prone mice. Proteinuria is a defining feature of glomerular filtration barrier dysfunction and a major clinical predictor of LN progression and long-term renal outcome [6–8,11,12,17–19]. In LN, persistent proteinuria may result from sustained immune complex-mediated injury involving complement activation and damage to podocytes, the glomerular endothelium, and the GBM [6–10,31–34]. In the present study, NP-treated mice maintained significantly lower and more stable protein-to-creatinine ratios than saline-treated LPR mice throughout disease progression, consistent with improved preservation of glomerular filtration barrier function. The reduction in proteinuria occurred alongside decreased histopathological injury and reduced glomerular IgG deposition, suggesting that the functional improvement was associated with structural renal protection rather than transient hemodynamic effects alone. reduction in proteinuria, GFR preservation was more modest, suggesting partial rather than complete protection of filtration capacity. The smaller terminal decline from baseline supports a treatment benefit, whereas the absence of a significant age × treatment interaction indicates that the overall longitudinal trajectories were not clearly separated. Several factors may contribute to this divergence. Proteinuria may improve before GFR because barrier injury can respond while pre-existing nephron loss and fibrosis remain incompletely reversible [6–8,11,12,17–19]. Measurement variability, attrition, and unequal survival may also reduce sensitivity in longitudinal GFR comparisons. Similar dissociations between proteinuria reduction and incomplete GFR recovery occur clinically during LN treatment [11,12,17–20]. The proteinuria findings therefore remain biologically relevant, but the GFR result should be presented as partial preservation rather than normalization

Histopathological analyses further demonstrated that targeted NP treatment markedly attenuated renal structural injury. Saline-treated MRL/lpr mice developed characteristic LN pathology, including mesangial expansion, glomerulosclerosis, capillary loop distortion, tubular injury, and inflammatory infiltrates [6–9,36–41]. These pathological changes were substantially reduced in NP-treated mice, as reflected by lower EGTI injury scores and reduced terminal kidney hypertrophy. These structural findings support the functional data by demonstrating that reduced proteinuria was accompanied by measurable attenuation of renal tissue injury, even though preservation of GFR was incomplete.

Current clinical treatment strategies for active LN commonly use pulse intravenous glucocorticoids followed by oral prednisone or prednisolone at doses up to approximately 0.5 mg/kg/day, with subsequent tapering toward lower maintenance doses [12, 15, 58]. In contrast, the present study used an average daily-equivalent prednisolone dose of 0.34 mg/kg/day in mice, corresponding by body surface area–based allometric scaling to an estimated human-equivalent dose of approximately 0.028 mg/kg/day 9 [56, 57]. Although this comparison should be interpreted cautiously because of species- and formulation-dependent differences in pharmacokinetics, metabolism, biodistribution, and drug release, the reduction in cumulative proteinuria burden and attenuation of terminal GFR decline at this comparatively low dose support the therapeutic rationale for targeted delivery. [56–57] Glomerular localization may increase the local therapeutic efficiency of prednisolone within the injured renal compartment without requiring high systemic glucocorticoid exposure. This is translationally relevant because, although glucocorticoids remain central to LN induction therapy, cumulative exposure contributes substantially to treatment-related morbidity in patients with SLE [11,12,15,16,25–28]. Strategies that maintain renal therapeutic efficacy while reducing systemic glucocorticoid exposure therefore remain highly desirable [25–30].

NP-delivered prednisolone was not associated with sustained blood-glucose elevation in either B6 or LPR mice. Because glucocorticoid-associated hyperglycemia contributes to treatment morbidity [25–28], this finding provides a preliminary glycemic-tolerability signal. However, blood glucose alone cannot establish reduced systemic glucocorticoid exposure or reduced metabolic toxicity. Dedicated pharmacokinetic, insulin-sensitivity, lipid, and longer-term endocrine assessments would be required to support those claims.

Consistent with the absence of sustained hyperglycemia, exploratory longitudinal DEXA measurements showed age-related increases in bone mineral content and density without detected treatment-associated differences in skeletal mineral accrual or body-fat percentage among the evaluated groups (Supplementary Figure 2). These findings did not reveal overt skeletal or body-composition abnormalities during the measured interval.

The prolonged survival of NP-treated LPR mice further indicates that the observed renal and anti-inflammatory effects translated into a meaningful disease-level benefit. Because mortality in MRL/lpr mice reflects the combined burden of systemic autoimmunity and progressive renal disease, survival represents an integrated measure of overall disease severity [36–41]. The survival benefit is therefore consistent with attenuation of disease progression. NP treatment also did not adversely affect body-weight trajectory or produce overt deterioration in B6 mice These findings also support preliminary tolerability but dedicated pharmacokinetic and systemic-safety studies are required before reduced off-target toxicity can be claimed.

The reduction in glomerular IgG deposition provides additional evidence that NP treatment attenuated immune complex–associated renal injury. Glomerular immune complexes are central to LN pathogenesis and promote complement activation, leukocyte recruitment, oxidative stress, and inflammatory mediator production [6–10,13,14]. However, C3d deposition did not differ significantly between saline- and NP-treated LPR mice, indicating that the lower IgG burden was not accompanied by a detectable reduction in terminal complement deposition. The reduction in IgG may therefore reflect decreased immune-complex accumulation or retention and attenuation of the inflammatory conditions that perpetuate glomerular injury, rather than generalized suppression of complement activity. Because prednisolone would not be expected to directly eliminate autoantibody production, localized delivery may instead reduce inflammatory amplification, glomerular permeability changes, and secondary immune-cell recruitment [6–10,13,14]. The accompanying reduction in dUTP/TUNEL-associated staining is consistent with lower DNA fragmentation and renal tissue injury.

The cytokine profiling data further support an anti-inflammatory effect of Col4-α3-targeted NP delivery. Saline-treated MRL/lpr mice exhibited increased renal abundance of multiple inflammatory and tissue-remodeling mediators, including CCL6/C10, IL-11, IL-28A/B, IL-7, IL-15, and platelet-derived endothelial cell growth factor (PD-ECGF) [13,14,47–53]. These mediators are associated with processes relevant to LN, including leukocyte recruitment, lymphocyte survival and activation, interferon-related immune regulation, fibrosis, angiogenesis, and tissue remodeling [13,14,47–53]. IL-7 and IL-15 are important regulators of T-cell survival, proliferation, and activation, whereas IL-11 has been associated with inflammatory and fibrotic responses in chronic tissue injury and has been reported as a urinary marker of disease activity [47–49]. IL-28A/B, also known as type III interferons or IFN-λ, contributes to innate and adaptive immune regulation and has been linked to inflammatory responses in autoimmune disease [50,51]. CCL6/C10 functions as a chemotactic mediator involved in leukocyte recruitment and has been reported to be elevated in murine LN kidneys [52], whereas PD-ECGF has been associated with angiogenesis and tissue remodeling in inflammatory environments [53]. The reduced abundance of these mediators following NP treatment is consistent with attenuation of inflammatory and tissue-remodeling processes within the renal microenvironment and complements our previous observations of reduced renal myeloid-cell and T-cell activation following Col4-α3-NP treatment [35].

Several limitations of the current study should be considered when interpreting these findings. Although the MRL/lpr model reproduces many important features of human LN, no single murine model captures the full immunologic and clinical heterogeneity of human SLE [36–41]. In addition, the main efficacy cohort did not include separate blank NP or free prednisolone comparator arms. These groups were not advanced into the definitive study because pilot experiments demonstrated similar longitudinal renal functional patterns and no significant difference in terminal relative kidney weight between blank NP- and dose-matched free prednisolone-treated MRL/lpr mice (Supplementary Figure 1). This pilot work supported prioritization of the targeted prednisolone-loaded NP and saline-control comparison in the primary cohort. Nevertheless, adequately powered comparator groups will be important for distinguishing the relative contributions of glomerular targeting, sustained drug release, systemic prednisolone exposure, and the NP carrier. The study also did not extensively characterize circulating autoantibodies, systemic immune responses, pharmacokinetics, or intrarenal biodistribution in diseased mice. Cytokine profiling did not establish cellular sources or pathway activation and should be interpreted as exploratory.

Future studies should evaluate the platform with additional therapeutic cargos and define how targeted prednisolone alters resident and infiltrating renal-cell populations. Flow cytometry, spatial profiling, transcriptomics, proteomics, and single-cell approaches could distinguish effects on immune-cell recruitment, activation, filtration-barrier integrity, and inflammatory-niche remodeling [54,55]. Further optimization of dosing and nanoparticle composition, together with comprehensive biodistribution, pharmacokinetic, and long-term safety assessment, will be required before clinical translation.

In conclusion, Col4-α3-Pred-NP delivery attenuated LN in MRL/lpr mice treated after disease onset. Treatment improved survival, reduced proteinuria burden and terminal GFR decline, decreased glomerular IgG deposition and renal structural injury, and reduced renal inflammatory-mediator abundance. Treatment was not associated with overt abnormalities in the measured glycemic, skeletal, body-composition, or body-weight outcomes during the evaluated interval. Together, these findings support glomerulus-targeted nanotherapy as a strategy with the potential to improve the precision and therapeutic index of glucocorticoid treatment in LN and other immune-mediated kidney diseases [11,12,15,16,25–34].

