## Supplementary Materials for "Glomerular-Targeted Delivery of Low-Dose Prednisolone Attenuates Established Lupus Nephritis in MRL/lpr Mice"

Supplementary Table S1. Animal-level survival status and study disposition of B6 and MRL/lpr mice following treatment.

*Animal identifiers M1-M40 correspond sequentially to Kaplan-Meier entries.*

| **Mouse ID** | **Experimental group** | **Last observed week (age)** | **Event code** | **Disposition** | **Reached week 20** |
| --- | --- | --- | --- | --- | --- |
| M1 | B6 Saline | 20 | 0 | Completed follow-up; censored at planned endpoint | Yes |
| M2 | B6 Saline | 20 | 0 | Completed follow-up; censored at planned endpoint | Yes |
| M3 | B6 Saline | 20 | 0 | Completed follow-up; censored at planned endpoint | Yes |
| M4 | B6 Saline | 20 | 0 | Completed follow-up; censored at planned endpoint | Yes |
| M5 | B6 Saline | 20 | 0 | Completed follow-up; censored at planned endpoint | Yes |
| M6 | B6 Saline | 14 | 0 | Early censored | No |
| M7 | B6 Saline | 20 | 0 | Completed follow-up; censored at planned endpoint | Yes |
| M8 | B6 Saline | 14 | 0 | Early censored | No |
| M9 | B6 Saline | 20 | 0 | Completed follow-up; censored at planned endpoint | Yes |
| M10 | B6 Saline | 14 | 0 | Early censored | No |
| M11 | B6 Col4-alpha3 NP | 20 | 0 | Completed follow-up; censored at planned endpoint | Yes |
| M12 | B6 Col4-alpha3 NP | 20 | 0 | Completed follow-up; censored at planned endpoint | Yes |
| M13 | B6 Col4-alpha3 NP | 20 | 0 | Completed follow-up; censored at planned endpoint | Yes |
| M14 | B6 Col4-alpha3 NP | 20 | 0 | Completed follow-up; censored at planned endpoint | Yes |
| M15 | B6 Col4-alpha3 NP | 20 | 0 | Completed follow-up; censored at planned endpoint | Yes |
| M16 | B6 Col4-alpha3 NP | 20 | 0 | Completed follow-up; censored at planned endpoint | Yes |
| M17 | B6 Col4-alpha3 NP | 20 | 0 | Completed follow-up; censored at planned endpoint | Yes |
| M18 | B6 Col4-alpha3 NP | 16 | 0 | Early censored | No |
| M19 | B6 Col4-alpha3 NP | 20 | 0 | Completed follow-up; censored at planned endpoint | Yes |
| M20 | B6 Col4-alpha3 NP | 20 | 0 | Completed follow-up; censored at planned endpoint | Yes |
| M21 | LPR Saline | 14 | 1 | Death event | No |
| M22 | LPR Saline | 18 | 1 | Death event | No |
| M23 | LPR Saline | 20 | 0 | Completed follow-up; censored at planned endpoint | Yes |
| M24 | LPR Saline | 20 | 0 | Completed follow-up; censored at planned endpoint | Yes |
| M25 | LPR Saline | 20 | 0 | Completed follow-up; censored at planned endpoint | Yes |
| M26 | LPR Saline | 15 | 1 | Death event | No |
| M27 | LPR Saline | 16 | 1 | Death event | No |
| M28 | LPR Saline | 20 | 0 | Completed follow-up; censored at planned endpoint | Yes |
| M29 | LPR Saline | 16 | 1 | Death event | No |
| M30 | LPR Saline | 16 | 1 | Death event | No |
| M31 | LPR Col4-alpha3 NP | 20 | 0 | Completed follow-up; censored at planned endpoint | Yes |
| M32 | LPR Col4-alpha3 NP | 20 | 0 | Completed follow-up; censored at planned endpoint | Yes |
| M33 | LPR Col4-alpha3 NP | 20 | 0 | Completed follow-up; censored at planned endpoint | Yes |
| M34 | LPR Col4-alpha3 NP | 20 | 0 | Completed follow-up; censored at planned endpoint | Yes |
| M35 | LPR Col4-alpha3 NP | 20 | 0 | Completed follow-up; censored at planned endpoint | Yes |
| M36 | LPR Col4-alpha3 NP | 20 | 0 | Completed follow-up; censored at planned endpoint | Yes |
| M37 | LPR Col4-alpha3 NP | 17 | 1 | Death event | No |
| M38 | LPR Col4-alpha3 NP | 17 | 1 | Death event | No |
| M39 | LPR Col4-alpha3 NP | 20 | 0 | Completed follow-up; censored at planned endpoint | Yes |
| M40 | LPR Col4-alpha3 NP | 20 | 0 | Completed follow-up; censored at planned endpoint | Yes |

**Notes:** Event code 1 denotes death, and event code 0 denotes censoring. Animals with code 0 at week 20 completed the planned follow-up and were censored at the study endpoint. Animals with code 0 before week 20 were censored early and contributed to the Kaplan-Meier analysis only through their last recorded week. “Reached week 20” indicates the maximum number potentially available for terminal procedures; exact assay-specific n values may be lower because of sample availability, prespecified exclusions, or technical quality-control failure.

| **Supplementary Table S2. Antibodies used for Immunofluorescence staining.** | | | | | | | |
| --- | --- | --- | --- | --- | --- | --- | --- |
| **Antibody** | **Target** | **Host** | **Fluorophore** | **Vendor** | **Catalog #** | **Application** | **Concentration** |
| Donkey anti-mouse IgG H&L | Endogenous mouse IgG immune-complex deposition | Donkey anti-mouse IgG | Alexa Fluor 488 | Abcam | ab150109 | IF-P | 1:1000 |
| Goat anti-mouse/rat complement component C3d | C3d / complement deposition | Goat polyclonal anti-mouse/rat C3d | Unconjugated primary antibody | R&D Systems / Bio-Techne | AF2655 | IF-P | 10ug/mL |
| Donkey anti-goat IgG H&L | Detection of goat anti-C3d primary antibody | Donkey anti-goat IgG | Alexa Fluor 594 | Abcam | ab150136 | IF-P | 1:1000 |

| Supplementary Table S3. Full cytokine array statistical summary for LPR saline versus LPR Col4-α3-NP-treated mice | | | | | | | | | | | |
| --- | --- | --- | --- | --- | --- | --- | --- | --- | --- | --- | --- |
| *Summary: 111 analytes tested; 6 nominal hits (6 down, 0 up); 0 BH/FDR-adjusted hits.* | | | | | | | | | | | |
| **Cytokine** | **Mean LPR Saline** | **Mean LPR Col4-α3-NP** | **Median LPR Saline** | **Median LPR Col4-α3-NP** | **Fold change (NP/Saline)** | **log2FC (NP vs Saline)** | **Nominal p value** | **BH-adjusted p value** | **Nominal hit** | **FDR hit** | **Regulation** |
| IL-28A/B | 0.0276 | 0.0002 | 0.0303 | 0.0001 | 0.0122 | -6.3528 | 0.0034 | 0.2191 | Yes | No | Down |
| IL-7 | 0.0515 | 0.0105 | 0.0511 | 0.0104 | 0.2055 | -2.2830 | 0.0039 | 0.2191 | Yes | No | Down |
| PD-ECGF | 0.0146 | 0.0017 | 0.0156 | 0.0008 | 0.1248 | -3.0022 | 0.0161 | 0.5911 | Yes | No | Down |
| IL-11 | 0.0393 | 0.0064 | 0.0373 | 0.0062 | 0.1649 | -2.6001 | 0.0216 | 0.5911 | Yes | No | Down |
| CCL6/C10 | 0.3932 | 0.0997 | 0.3995 | 0.0689 | 0.2537 | -1.9789 | 0.0349 | 0.5911 | Yes | No | Down |
| IL-15 | 0.0274 | 0.0006 | 0.0292 | 0.0000 | 0.0269 | -5.2182 | 0.0411 | 0.5911 | Yes | No | Down |
| Flt-3 Ligand | 0.0982 | 0.0176 | 0.1101 | 0.0073 | 0.1796 | -2.4773 | 0.0568 | 0.5911 | No | No | Not significant |
| Pref-1/DLK-1/FA1 | 0.0155 | 0.0008 | 0.0155 | 0.0000 | 0.0578 | -4.1129 | 0.0577 | 0.5911 | No | No | Not significant |
| IL-12 p40 | 0.0645 | 0.0087 | 0.0689 | 0.0038 | 0.1354 | -2.8847 | 0.0606 | 0.5911 | No | No | Not significant |
| M-CSF | 0.0715 | 0.0186 | 0.0632 | 0.0156 | 0.2606 | -1.9404 | 0.0815 | 0.5911 | No | No | Not significant |
| Pentraxin 3/TSG-14 | 0.0203 | 0.0059 | 0.0205 | 0.0051 | 0.2961 | -1.7557 | 0.0850 | 0.5911 | No | No | Not significant |
| IL-1a/IL-1F1 | 0.0324 | 0.0012 | 0.0345 | 0.0000 | 0.0387 | -4.6914 | 0.0863 | 0.5911 | No | No | Not significant |
| CXCL13/BLC/BCA-1 | 0.2194 | 0.0063 | 0.1691 | 0.0000 | 0.0291 | -5.1015 | 0.0924 | 0.5911 | No | No | Not significant |
| IL-13 | 0.0139 | 0.0021 | 0.0161 | 0.0000 | 0.1599 | -2.6445 | 0.0949 | 0.5911 | No | No | Not significant |
| DPPIV/CD26 | 0.2342 | 0.1416 | 0.2298 | 0.1358 | 0.6047 | -0.7256 | 0.1044 | 0.5911 | No | No | Not significant |
| Serpin F1/PEDF | 0.0292 | 0.0135 | 0.0296 | 0.0109 | 0.4619 | -1.1145 | 0.1065 | 0.5911 | No | No | Not significant |
| Reg3G | 0.5799 | 0.3851 | 0.5584 | 0.4191 | 0.6641 | -0.5905 | 0.1067 | 0.5911 | No | No | Not significant |
| RBP4 | 0.3353 | 0.2238 | 0.3397 | 0.2183 | 0.6676 | -0.5829 | 0.1205 | 0.5911 | No | No | Not significant |
| IFN-y | 0.0585 | 0.0084 | 0.0601 | 0.0000 | 0.1452 | -2.7838 | 0.1220 | 0.5911 | No | No | Not significant |
| GM-CSF | 0.0142 | 0.0000 | 0.0123 | 0.0000 | 0.0063 | -7.3030 | 0.1232 | 0.5911 | No | No | Not significant |
| HGF | 0.1224 | 0.0413 | 0.1373 | 0.0081 | 0.3378 | -1.5657 | 0.1248 | 0.5911 | No | No | Not significant |
| DKK-1 | 0.0261 | 0.0000 | 0.0224 | 0.0000 | 0.0034 | -8.1801 | 0.1264 | 0.5911 | No | No | Not significant |
| Proliferin | 0.0172 | 0.0000 | 0.0169 | 0.0000 | 0.0052 | -7.5794 | 0.1325 | 0.5911 | No | No | Not significant |
| IL-5 | 0.0129 | 0.0023 | 0.0131 | 0.0000 | 0.1846 | -2.4375 | 0.1330 | 0.5911 | No | No | Not significant |
| Osteoprotegerin/TNFRSF11B | 0.0212 | 0.0065 | 0.0252 | 0.0041 | 0.3077 | -1.7003 | 0.1374 | 0.5911 | No | No | Not significant |
| IL-10 | 0.0410 | 0.0172 | 0.0318 | 0.0163 | 0.4219 | -1.2449 | 0.1385 | 0.5911 | No | No | Not significant |
| Osteopontin (OPN) | 0.3307 | 0.2162 | 0.3493 | 0.1876 | 0.6537 | -0.6133 | 0.1463 | 0.6016 | No | No | Not significant |
| Resistin | 0.3272 | 0.2168 | 0.3197 | 0.2145 | 0.6626 | -0.5938 | 0.1613 | 0.6212 | No | No | Not significant |
| Myeloperoxidase | 0.1726 | 0.1051 | 0.1691 | 0.1065 | 0.6091 | -0.7153 | 0.1623 | 0.6212 | No | No | Not significant |
| IL-6 | 0.0392 | 0.0032 | 0.0304 | 0.0000 | 0.0825 | -3.6002 | 0.1727 | 0.6391 | No | No | Not significant |
| Cystatin C | 0.3978 | 0.3046 | 0.4047 | 0.2880 | 0.7656 | -0.3853 | 0.1823 | 0.6482 | No | No | Not significant |
| IL-3 | 0.0151 | 0.0024 | 0.0134 | 0.0000 | 0.1621 | -2.6250 | 0.1869 | 0.6482 | No | No | Not significant |
| Adiponectin/Acrp30 | 0.1260 | 0.0549 | 0.1145 | 0.0440 | 0.4358 | -1.1982 | 0.2059 | 0.6532 | No | No | Not significant |
| BAFF/BLyS/TNFSF13B | 0.0935 | 0.0335 | 0.1005 | 0.0186 | 0.3587 | -1.4793 | 0.2133 | 0.6532 | No | No | Not significant |
| LIX | 0.0164 | 0.0029 | 0.0126 | 0.0000 | 0.1818 | -2.4595 | 0.2209 | 0.6532 | No | No | Not significant |
| RAGE | 0.0418 | 0.0138 | 0.0328 | 0.0061 | 0.3313 | -1.5937 | 0.2271 | 0.6532 | No | No | Not significant |
| IL-4 | 0.0831 | 0.0069 | 0.0437 | 0.0000 | 0.0842 | -3.5698 | 0.2272 | 0.6532 | No | No | Not significant |
| IL-27 p28 | 0.0245 | 0.0113 | 0.0265 | 0.0101 | 0.4630 | -1.1111 | 0.2281 | 0.6532 | No | No | Not significant |
| C-Reactive Protein/CRP | 0.1215 | 0.0577 | 0.1292 | 0.0415 | 0.4751 | -1.0738 | 0.2373 | 0.6532 | No | No | Not significant |
| CXCL11/1-TAC | 0.0316 | 0.0042 | 0.0203 | 0.0000 | 0.1354 | -2.8844 | 0.2401 | 0.6532 | No | No | Not significant |
| Gas 6 | 0.1550 | 0.0557 | 0.1517 | 0.0112 | 0.3600 | -1.4741 | 0.2413 | 0.6532 | No | No | Not significant |
| FGF-21 | 0.0253 | 0.0025 | 0.0115 | 0.0000 | 0.1034 | -3.2739 | 0.2535 | 0.6700 | No | No | Not significant |
| IL-17A | 0.0060 | 0.0004 | 0.0033 | 0.0000 | 0.0747 | -3.7426 | 0.2623 | 0.6722 | No | No | Not significant |
| LIF | 0.0410 | 0.0179 | 0.0345 | 0.0188 | 0.4371 | -1.1939 | 0.2665 | 0.6722 | No | No | Not significant |
| Serpin E1/PAI-1 | 0.1412 | 0.0772 | 0.1711 | 0.0680 | 0.5467 | -0.8711 | 0.2911 | 0.6986 | No | No | Not significant |
| IGFBP-1 | 0.4665 | 0.0262 | 0.1918 | 0.0000 | 0.0564 | -4.1488 | 0.2914 | 0.6986 | No | No | Not significant |
| Lipocalin-2/NGAL | 0.2077 | 0.1330 | 0.2086 | 0.1205 | 0.6406 | -0.6425 | 0.3005 | 0.6986 | No | No | Not significant |
| Complement Component C5/C5a | 0.0327 | 0.0133 | 0.0344 | 0.0035 | 0.4067 | -1.2979 | 0.3053 | 0.6986 | No | No | Not significant |
| MMP-3 | 0.1203 | 0.0504 | 0.0869 | 0.0357 | 0.4193 | -1.2539 | 0.3199 | 0.6986 | No | No | Not significant |
| GDF-15 | 0.0313 | 0.0128 | 0.0361 | 0.0000 | 0.4100 | -1.2865 | 0.3268 | 0.6986 | No | No | Not significant |
| IL-23 | 0.0108 | 0.0037 | 0.0110 | 0.0000 | 0.3482 | -1.5220 | 0.3304 | 0.6986 | No | No | Not significant |
| TNF-a | 0.0044 | 0.0014 | 0.0037 | 0.0009 | 0.3273 | -1.6113 | 0.3367 | 0.6986 | No | No | Not significant |
| WISP-1/CCN4 | 0.0360 | 0.0299 | 0.0329 | 0.0277 | 0.8309 | -0.2672 | 0.3389 | 0.6986 | No | No | Not significant |
| MMP-9 | 0.0169 | 0.0053 | 0.0118 | 0.0000 | 0.3183 | -1.6515 | 0.3400 | 0.6986 | No | No | Not significant |
| IL-1B/IL-1F2 | 0.0141 | 0.0047 | 0.0113 | 0.0000 | 0.3378 | -1.5657 | 0.3461 | 0.6986 | No | No | Not significant |
| CD14 | 0.1032 | 0.0521 | 0.1077 | 0.0467 | 0.5050 | -0.9855 | 0.3542 | 0.7020 | No | No | Not significant |
| FGF acidic | 0.6618 | 0.7964 | 0.6540 | 0.8715 | 1.2034 | 0.2671 | 0.3619 | 0.7047 | No | No | Not significant |
| CCL20/MIP-3a | 0.0264 | 0.3205 | 0.0230 | 0.0635 | 12.1161 | 3.5988 | 0.3687 | 0.7055 | No | No | Not significant |
| P-Selectin/CD62P | 0.0510 | 0.0320 | 0.0588 | 0.0242 | 0.6267 | -0.6742 | 0.3897 | 0.7193 | No | No | Not significant |
| IL-22 | 0.0066 | 0.0007 | 0.0011 | 0.0000 | 0.1193 | -3.0669 | 0.3915 | 0.7193 | No | No | Not significant |
| Chemerin | 0.0861 | 0.0507 | 0.0997 | 0.0433 | 0.5887 | -0.7645 | 0.3953 | 0.7193 | No | No | Not significant |
| CXCL10/IP-10 | 0.0257 | 0.0061 | 0.0090 | 0.0000 | 0.2397 | -2.0609 | 0.4039 | 0.7231 | No | No | Not significant |
| MMP-2 | 0.1112 | 0.0357 | 0.0555 | 0.0010 | 0.3218 | -1.6356 | 0.4239 | 0.7458 | No | No | Not significant |
| G-CSF | 0.0293 | 0.0144 | 0.0356 | 0.0000 | 0.4941 | -1.0170 | 0.4350 | 0.7458 | No | No | Not significant |
| CCL3/CCL4/MIP-1a/ß | 0.0357 | 0.0167 | 0.0324 | 0.0000 | 0.4678 | -1.0961 | 0.4423 | 0.7458 | No | No | Not significant |
| Thrombopoietin | 0.0067 | 0.0027 | 0.0052 | 0.0000 | 0.4131 | -1.2755 | 0.4434 | 0.7458 | No | No | Not significant |
| IL-2 | 0.0027 | 0.0004 | 0.0000 | 0.0000 | 0.1805 | -2.4699 | 0.4616 | 0.7647 | No | No | Not significant |
| Periostin/OSF-2 | 0.0095 | 0.0047 | 0.0090 | 0.0000 | 0.4942 | -1.0169 | 0.4766 | 0.7779 | No | No | Not significant |
| Coagulation Factor III/Tissue Factor | 0.3277 | 0.1957 | 0.2977 | 0.1088 | 0.5972 | -0.7438 | 0.4918 | 0.7835 | No | No | Not significant |
| CCL17/TARC | 0.0369 | 0.0197 | 0.0303 | 0.0080 | 0.5362 | -0.8992 | 0.4941 | 0.7835 | No | No | Not significant |
| CCL22/MDC | 0.0454 | 0.0255 | 0.0460 | 0.0030 | 0.5620 | -0.8312 | 0.5330 | 0.8302 | No | No | Not significant |
| VCAM-1/CD106 | 0.5127 | 0.3986 | 0.5021 | 0.3518 | 0.7773 | -0.3634 | 0.5397 | 0.8302 | No | No | Not significant |
| IGFBP-5 | 0.1640 | 0.1136 | 0.1649 | 0.0678 | 0.6925 | -0.5301 | 0.5474 | 0.8302 | No | No | Not significant |
| IL-1ra/IL-1F3 | 0.1109 | 0.0619 | 0.0909 | 0.0112 | 0.5579 | -0.8418 | 0.5535 | 0.8302 | No | No | Not significant |
| Amphiregulin | 0.0291 | 0.0128 | 0.0085 | 0.0000 | 0.4406 | -1.1824 | 0.5738 | 0.8383 | No | No | Not significant |
| IGFBP-2 | 0.2639 | 0.1852 | 0.2135 | 0.1536 | 0.7019 | -0.5107 | 0.5822 | 0.8383 | No | No | Not significant |
| IGFBP-6 | 0.3514 | 0.2707 | 0.3154 | 0.2793 | 0.7706 | -0.3760 | 0.5899 | 0.8383 | No | No | Not significant |
| Leptin | 0.0207 | 0.0124 | 0.0122 | 0.0113 | 0.5997 | -0.7376 | 0.5905 | 0.8383 | No | No | Not significant |
| Fetuin A/AHSG | 0.3099 | 0.2636 | 0.3166 | 0.2587 | 0.8507 | -0.2333 | 0.6097 | 0.8383 | No | No | Not significant |
| Angiopoietin-2 | 0.2486 | 0.1684 | 0.2382 | 0.1009 | 0.6776 | -0.5615 | 0.6118 | 0.8383 | No | No | Not significant |
| Endoglin/CD105 | 0.4436 | 0.3785 | 0.4498 | 0.3653 | 0.8534 | -0.2287 | 0.6277 | 0.8383 | No | No | Not significant |
| IGFBP-3 | 0.1691 | 0.1449 | 0.1636 | 0.1691 | 0.8565 | -0.2235 | 0.6291 | 0.8383 | No | No | Not significant |
| CCL2/JE/MCP-1 | 0.0349 | 0.0218 | 0.0352 | 0.0000 | 0.6251 | -0.6780 | 0.6342 | 0.8383 | No | No | Not significant |
| PDGF-BB | 0.0478 | 0.0414 | 0.0456 | 0.0466 | 0.8663 | -0.2071 | 0.6421 | 0.8383 | No | No | Not significant |
| C1q R1/CD93 | 0.3972 | 0.2934 | 0.4085 | 0.2448 | 0.7389 | -0.4365 | 0.6535 | 0.8383 | No | No | Not significant |
| E-Selectin/CD62E | 0.0552 | 0.0661 | 0.0500 | 0.0495 | 1.1963 | 0.2586 | 0.6608 | 0.8383 | No | No | Not significant |
| Chitinase 3-like 1 | 0.4778 | 0.4148 | 0.4683 | 0.5029 | 0.8682 | -0.2039 | 0.6766 | 0.8383 | No | No | Not significant |
| Angiopoietin-1 | 0.0374 | 0.0261 | 0.0323 | 0.0124 | 0.6977 | -0.5194 | 0.6895 | 0.8383 | No | No | Not significant |
| Complement Factor D | 0.0661 | 0.0512 | 0.0759 | 0.0410 | 0.7741 | -0.3695 | 0.7146 | 0.8383 | No | No | Not significant |
| CD40/TNFRSFS | 0.4507 | 0.3716 | 0.4909 | 0.2947 | 0.8246 | -0.2782 | 0.7184 | 0.8383 | No | No | Not significant |
| LDL R | 0.1668 | 0.1370 | 0.1432 | 0.1019 | 0.8213 | -0.2840 | 0.7206 | 0.8383 | No | No | Not significant |
| EGF | 0.4627 | 0.3866 | 0.4875 | 0.3610 | 0.8357 | -0.2590 | 0.7240 | 0.8383 | No | No | Not significant |
| CCL11/Eotaxin | 0.0313 | 0.0204 | 0.0097 | 0.0070 | 0.6524 | -0.6163 | 0.7278 | 0.8383 | No | No | Not significant |
| TIM-1/KIM-1/HAVCR | 0.2695 | 0.3125 | 0.2684 | 0.2854 | 1.1598 | 0.2139 | 0.7282 | 0.8383 | No | No | Not significant |
| CXCL2/MIP-2 | 0.0397 | 0.0288 | 0.0405 | 0.0064 | 0.7264 | -0.4611 | 0.7343 | 0.8383 | No | No | Not significant |
| CXCL9/MIG | 0.2292 | 0.1786 | 0.1755 | 0.1852 | 0.7792 | -0.3599 | 0.7379 | 0.8383 | No | No | Not significant |
| Proprotein Convertase 9/PCSK9 | 0.0960 | 0.1320 | 0.0839 | 0.0521 | 1.3740 | 0.4584 | 0.7414 | 0.8383 | No | No | Not significant |
| Angiopoietin-like 3 | 0.5359 | 0.4882 | 0.5275 | 0.5475 | 0.9110 | -0.1345 | 0.7455 | 0.8383 | No | No | Not significant |
| CD160 | 0.0258 | 0.0171 | 0.0103 | 0.0000 | 0.6662 | -0.5860 | 0.7494 | 0.8383 | No | No | Not significant |
| CCL19/MIP-3B | 0.0369 | 0.0279 | 0.0351 | 0.0122 | 0.7581 | -0.3996 | 0.7552 | 0.8383 | No | No | Not significant |
| CX3CL1/Fractalkine | 0.6960 | 0.7829 | 0.6727 | 0.6172 | 1.1248 | 0.1697 | 0.8202 | 0.8862 | No | No | Not significant |
| IL-33 | 0.6008 | 0.5446 | 0.5672 | 0.5006 | 0.9066 | -0.1415 | 0.8257 | 0.8862 | No | No | Not significant |
| VEGF | 0.0429 | 0.0373 | 0.0416 | 0.0263 | 0.8705 | -0.2001 | 0.8292 | 0.8862 | No | No | Not significant |
| CXCL1/KC | 0.0408 | 0.0346 | 0.0495 | 0.0198 | 0.8477 | -0.2384 | 0.8303 | 0.8862 | No | No | Not significant |
| CCL21/6Ckine | 1.5259 | 1.3715 | 1.5061 | 1.3061 | 0.8988 | -0.1539 | 0.8448 | 0.8930 | No | No | Not significant |
| Endostatin | 0.5056 | 0.5193 | 0.5015 | 0.5098 | 1.0271 | 0.0386 | 0.8576 | 0.8980 | No | No | Not significant |
| CCL12/MCP-5 | 0.0526 | 0.0595 | 0.0419 | 0.0468 | 1.1303 | 0.1768 | 0.8772 | 0.9100 | No | No | Not significant |
| CXCL16 | 0.5846 | 0.5601 | 0.5356 | 0.6576 | 0.9580 | -0.0619 | 0.8941 | 0.9189 | No | No | Not significant |
| CCL5/RANTES | 0.2562 | 0.2377 | 0.2411 | 0.2256 | 0.9279 | -0.1080 | 0.9116 | 0.9236 | No | No | Not significant |
| Pentraxin 2/SAP | 1.0538 | 1.0853 | 0.9856 | 1.0360 | 1.0299 | 0.0424 | 0.9153 | 0.9236 | No | No | Not significant |
| ICAM-1/CD54 | 0.7568 | 0.7557 | 0.7047 | 0.6916 | 0.9986 | -0.0021 | 0.9965 | 0.9965 | No | No | Not significant |
| *Footnote: Normalized cytokine array values were compared between LPR saline and LPR Col4-α3-NP-treated mice. Fold change was calculated as Col4-α3-NP divided by saline. P values represent nominal comparisons, and Benjamini-Hochberg-adjusted p values are provided for false-discovery-rate transparency. Regulation labels are based on nominal p < 0.05 and \|log2 fold change\| ≥ 0.5. No analytes met BH/FDR-adjusted significance.* | | | | | | | | | | | |

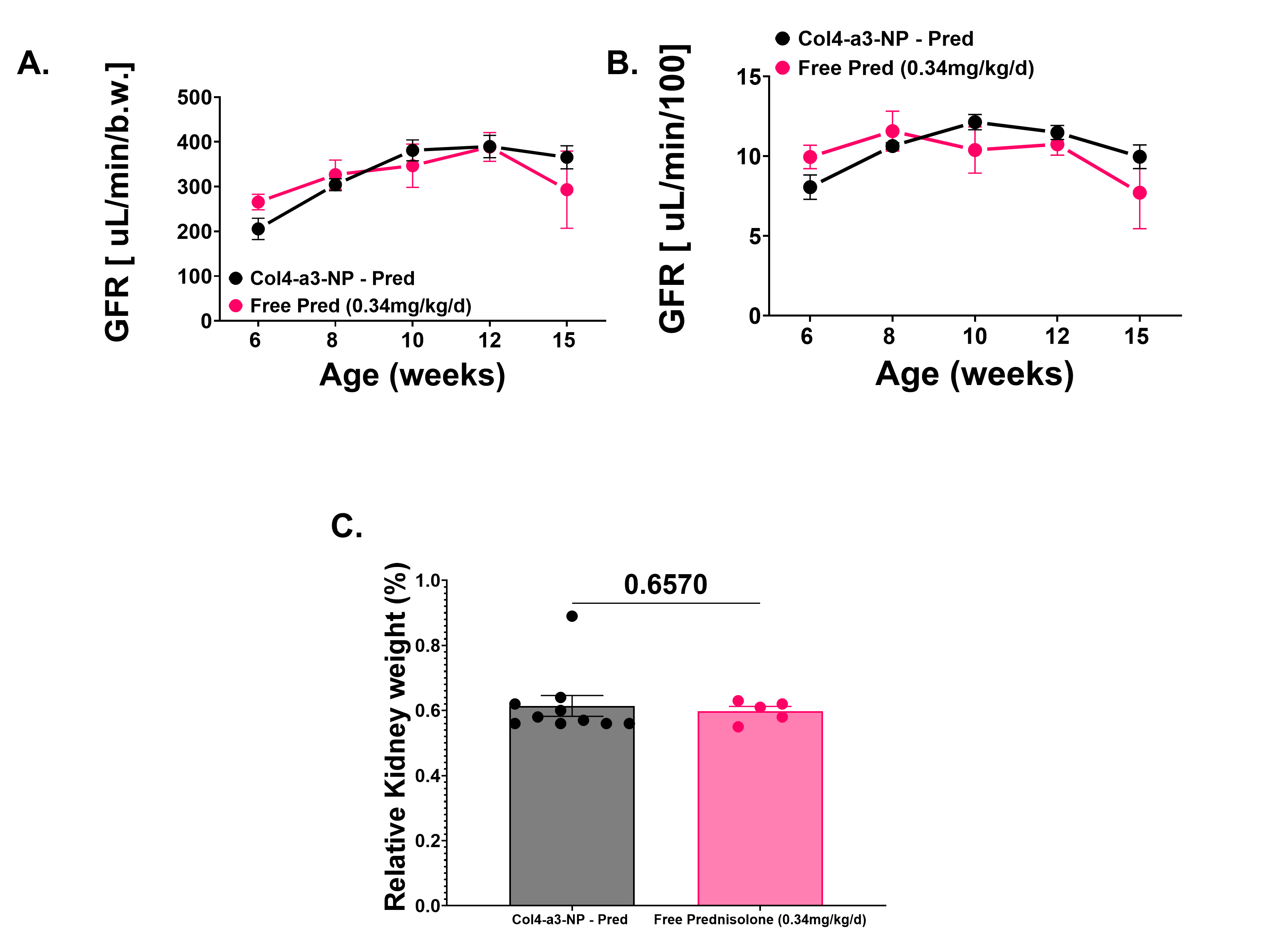

**Supplementary Figure 1. Free prednisolone and blank nanoparticles show comparable effects on renal function and kidney weight in lupus-prone mice.**
**A.** Longitudinal glomerular filtration rate (GFR) measured from 6 to 15 weeks of age in MRL/lpr mice treated with blank nanoparticles or free prednisolone. **B.** Age-stratified GFR values normalized to body weight across the study period. Ordinary two-way ANOVA demonstrated a significant effect of age/row factor on GFR normalized to body weight, indicating that renal filtration changed over time (p = 0.0036). However, there was no significant treatment effect between blank NP and free prednisolone groups (p = 0.8364) and no significant age × treatment interaction (p = 0.4870), indicating that the longitudinal GFR pattern was comparable between groups. **C.** Relative kidney weight at endpoint was not significantly different between blank nanoparticle-treated mice and mice treated with free prednisolone at 0.34 mg/kg/day (p = 0.6570). Individual points represent biological replicates, and bars show group mean ± SEM.

**Supplementary Figure 2. Prednisolone-loaded nanoparticles and free prednisolone do not adversely affect skeletal mineral accrual, body composition, or body-weight trajectories.**

**A.** Longitudinal bone mineral content measured by dual-energy X-ray absorptiometry at 6, 13, and 15 weeks of age in mice treated with blank Col4-α3 nanoparticles, prednisolone-loaded Col4-α3 nanoparticles at 0.34 mg/kg/day, or free prednisolone at 0.34 mg/kg/day. The 6-week measurement was obtained before treatment initiation and served as the baseline. Mixed-effects analysis demonstrated a significant effect of age on bone mineral content (p < 0.0001), indicating increased skeletal mineral accrual over time. However, there was no significant treatment effect (p = 0.6484) and no significant age × treatment interaction (p = 0.4033), indicating comparable longitudinal patterns among groups. B. Bone mineral density increased significantly with age (p = 0.0001), with no significant treatment effect (p = 0.9517) or age × treatment interaction (p = 0.4318). C. Body fat percentage was not significantly affected by age (p = 0.2218), treatment (p = 0.4213), or the age × treatment interaction (p = 0.3209). D. Longitudinal body-weight measurements from 6 to 17 weeks of age demonstrated a significant effect of age (p < 0.0001), whereas the treatment effect (p = 0.3673) and age × treatment interaction (p = 0.3351) were not significant, indicating comparable body-weight trajectories among groups. Individual points represent biological replicates, and body-weight data are shown as group mean ± SEM. n = 5, with one missing observation.

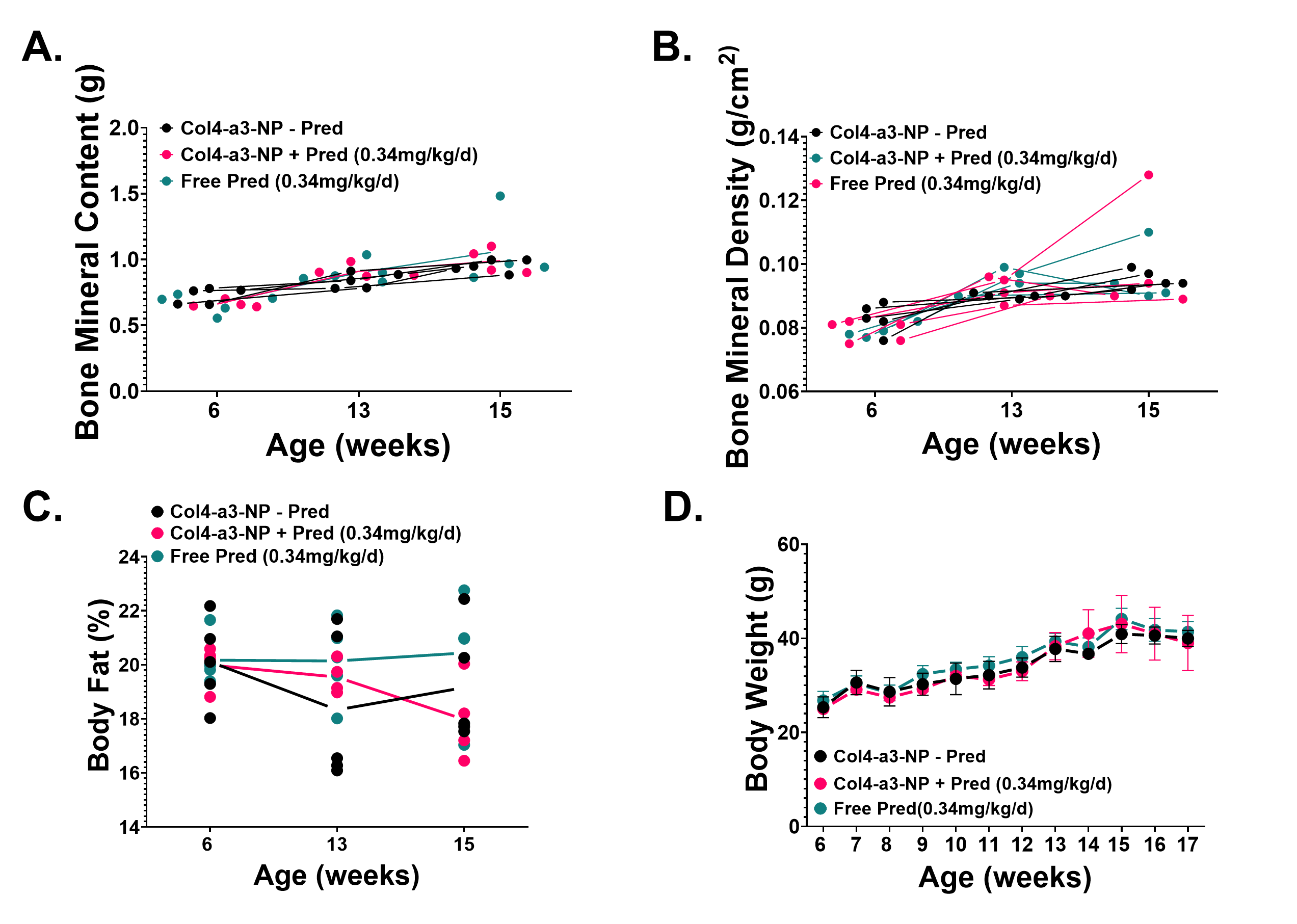

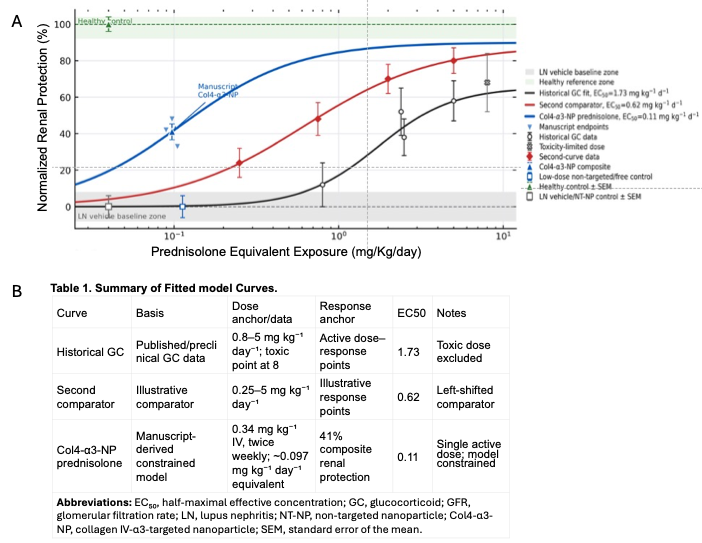

**Supplementary Figure 3. Comparative dose–response modeling of prednisolone-equivalent renal protection in preclinical LN studies. A.** Historical glucocorticoid dose–response data from preclinical LN studies were curated, digitized, normalized to study- specific vehicle controls, and compared with the experimentally observed Col4-α3-Pred-NP treatment point. The Col4-α3–NP curve represents a constrained pharmacodynamic model anchored to a single experimentally tested active dose and should not be interpreted as a fully fitted dose–response curve. Healthy control and MRL/lpr vehicle reference groups are shown as physiological benchmarks. EC50 values are model-derived estimates for comparative visualization only. Statistical and modeling assumptions are summarized in the accompanying table.

**Supplementary Figure 4. Col4-α3-NP treatment does not significantly alter renal cytokine expression in B6 control mice.**
**A.** Quantification of cytokine array targets in B6 saline and B6 Col4-α3-NP-treated mice. Individual cytokines were normalized and compared between groups; no cytokines were significantly altered by Col4-α3-NP treatment. Individual points represent biological replicates, and boxplots show the distribution of normalized values for each group. **B.** Row z-score heatmap of B6 renal cytokine profiles showing relative cytokine abundance across individual B6 saline and B6 Col4-α3-NP-treated mice. Columns represent individual mice and rows represent cytokine targets. Overall clustering and signal distribution did not reveal a consistent treatment-associated inflammatory shift in B6 kidneys.

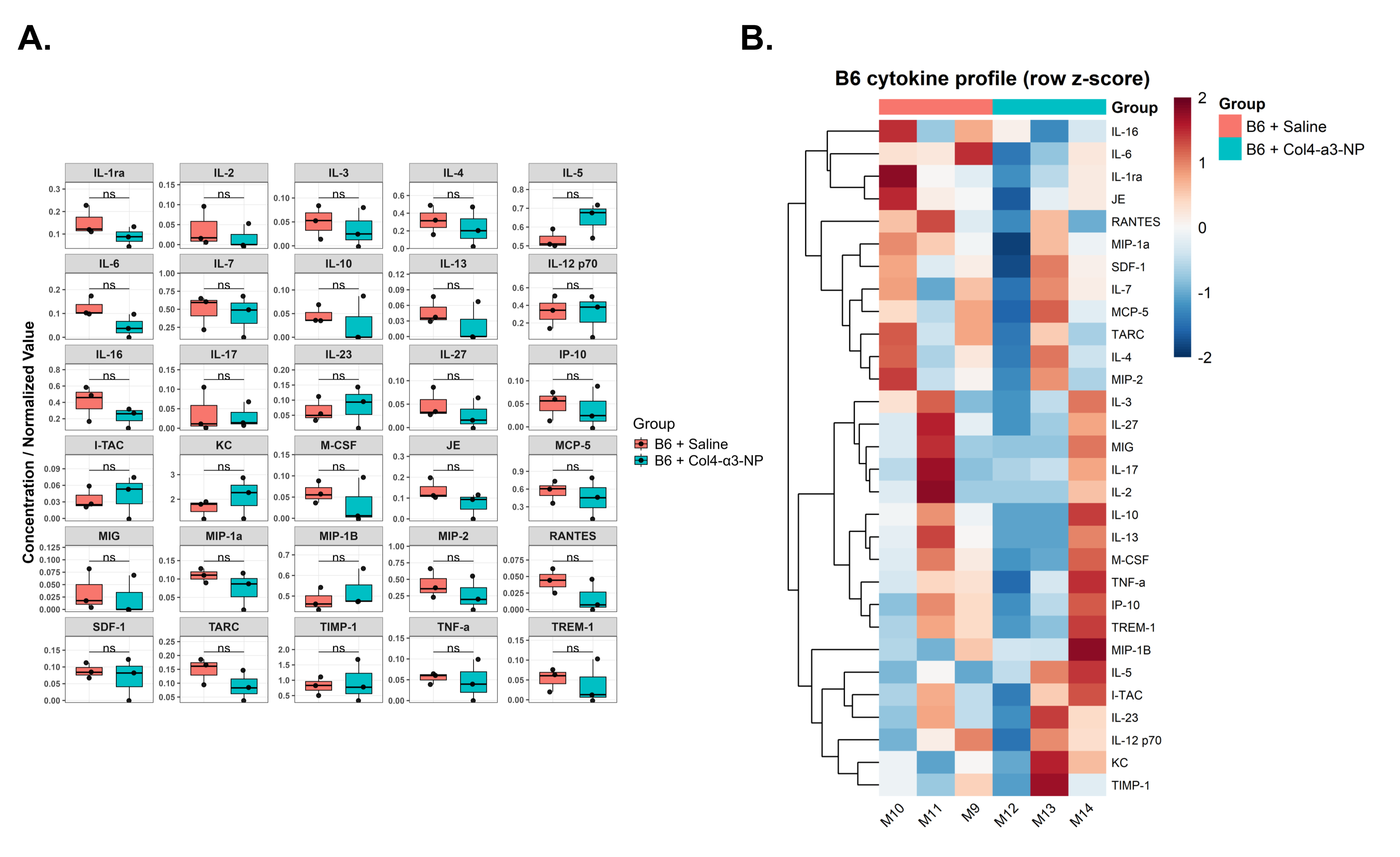

**Supplementary Figure 5. Normalized spleen weight is increased in MRL/lpr mice but is not significantly altered by Col4-α3-NP treatment.**
Terminal normalized spleen weight was assessed in B6 and MRL/lpr mice treated with saline or Col4-α3-NP. LPR mice displayed significantly increased relative spleen weight compared with B6 controls, consistent with lupus-associated splenomegaly. Col4-α3-NP treatment did not significantly alter normalized spleen weight in LPR mice compared with LPR saline controls (p = 0.59). Individual points represent biological replicates, and bars show group mean ± SEM. Statistical comparisons were performed using two-way ANOVA with post hoc multiple-comparison testing.

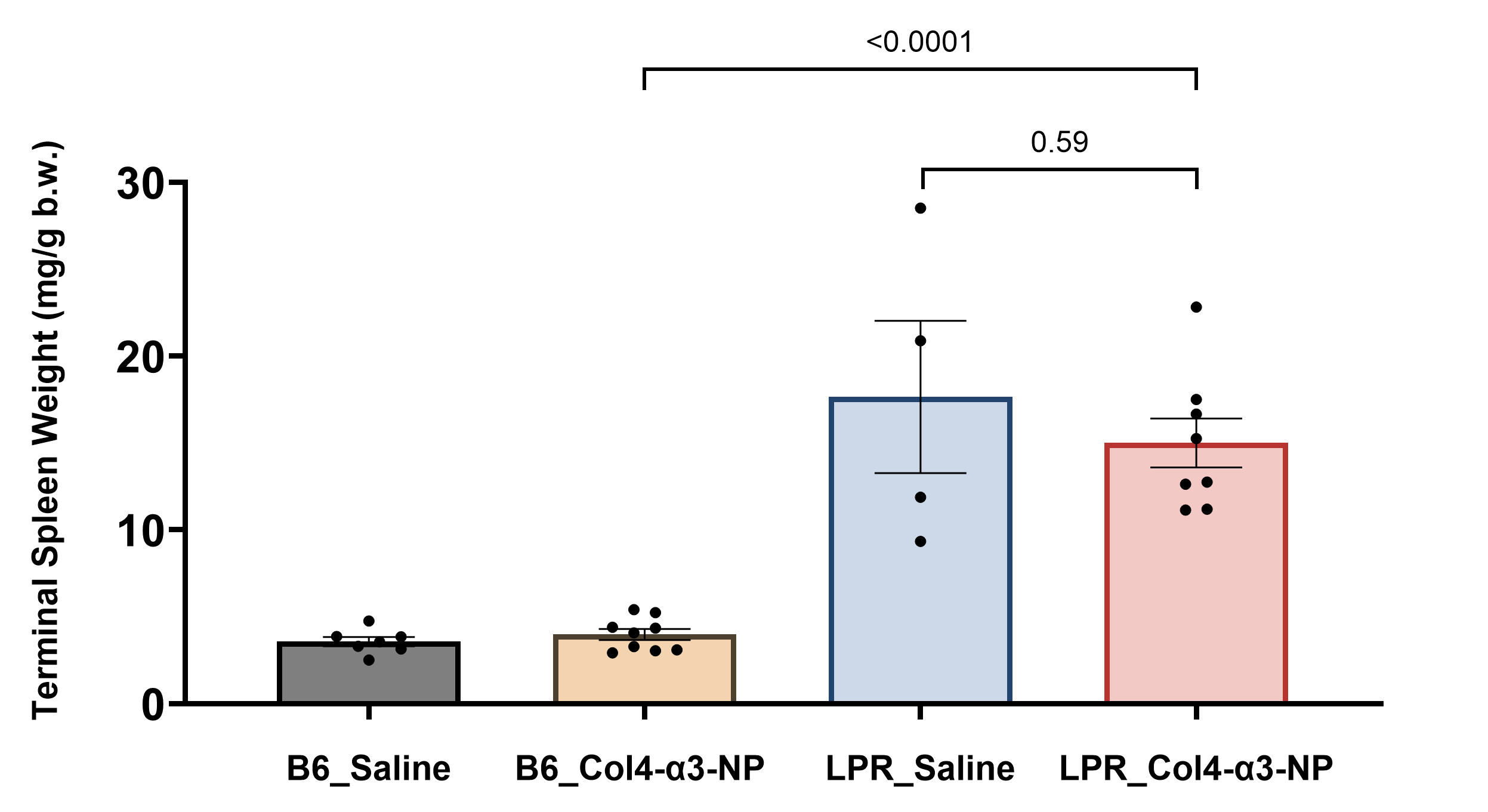
